# *Listeria monocytogenes* infection in pregnant macaques alters the maternal gut microbiome

**DOI:** 10.1101/2023.06.18.545418

**Authors:** Anna Marie Hugon, Courtney L. Deblois, Heather A. Simmons, Andres Mejia, Michele L. Schotzo, Charles J. Czuprynski, Garret Suen, Thaddeus G. Golos

## Abstract

**Objectives:** The bacterium *Listeria monocytogenes* (Lm) is associated with adverse pregnancy outcomes. Infection occurs through consumption of contaminated food that is disseminated to the maternal-fetal interface. The influence on the gastrointestinal microbiome during Lm infection remains unexplored in pregnancy. The objective of this study was to determine the impact of listeriosis on the gut microbiota of pregnant macaques.

**Methods:** A nonhuman primate model of listeriosis in pregnancy has been previously described [1, 2]. Both pregnant and nonpregnant cynomolgus macaques were inoculated with L. monocytogenes and bacteremia and fecal shedding were monitored for 14 days. Nonpregnant animal tissues were collected at necropsy to determine bacterial burden, and fecal samples from both pregnant and nonpregnant animals were evaluated by 16S rRNA next-generation sequencing.

**Results:** Unlike pregnant macaques, nonpregnant macaques did not exhibit bacteremia, fecal shedding, or tissue colonization by Lm. Dispersion of Lm during pregnancy was associated with a significant decrease in alpha-diversity of the host gut microbiome, compared to nonpregnant counterparts. The combined effects of pregnancy and listeriosis were associated with a significant loss in microbial richness, although there were increases in some genera and decreases in others.

**Conclusions:** Although pregnancy alone is not associated with gut microbiome disruption, we observed dysbiosis with listeriosis during pregnancy. The macaque model may provide an understanding of the roles that pregnancy and the gut microbiota play in the ability of Lm to establish intestinal infection and disseminate throughout the host, thereby contributing to adverse pregnancy outcomes and risk to the developing fetus.

**Summary sentence:** Intestinal microbial composition in macaques is influenced by significant interaction between the pregnant state and exposure to *Listeria monocytogenes*, associated in particular with significant changes to *Akkermansia, Eubacteria ruminantum, Methanobrevibacter, Prevotella,* and *Treponema*.

## Introduction

*Listeria monocytogenes* (Lm) is bacterial pathogen associated with fever, muscle aches, gastrointestinal upset, sepsis, and meningitis. It is a ubiquitous bacterium found in the environment and infection with Lm occurs via consumption of contaminated food. It has been shown that at-risk groups for listeriosis include young children, the elderly people, immunocompromised individuals, and pregnant women and their neonates. During pregnancy, infection can lead to serious complications including miscarriage, stillbirth, preterm birth, neonatal sepsis, and meningitis. Pregnant women infected with Lm are typically asymptomatic, lacking clinical features common in infected elderly or immunocompromised individuals [3]. As a result, maternal listeriosis may go unrecognized until infection of the maternal-fetal interface (MFI) resulting in adverse pregnancy outcomes (APOs). Importantly, although Lm does not cause severe illness or pathology within the mother, it is able to establish significant bacterial burden within the placenta and decidua, leading to severe infection, acute inflammation, and severe disruption to the MFI.

Although listeriosis can be treated with antibiotics such as ampicillin or gentamicin, these are only effective if diagnosis and administration occurs early during infection[4]. There has been growing concern over antimicrobial resistance in pathogens and the ability to survive at clinical antibiotic concentrations. Antibiotic resistance of Lm has been most notable in isolates from food products[5, 6], however rising rates of antibiotic resistance in humans from low-income countries is a concern for successful treatment and therapies of listeriosis [4, 7]. Moreover, antibiotics can have short term and long-term side effects such as gastrointestinal upset and neurotoxicity [8]. These observations provide additional impetus for alternatives to antibiotic treatment for listeriosis.

Previous research has sought to understand the molecular mechanisms behind the pathogenic ability of Lm to infect the placenta, with a focus on factors such as internalins and host immune interactions [9–12]. However, few studies focus on the maternal gut microbiome during infection with a bacterial pathogen [13–17]. From birth, the human gastrointestinal tract accumulates and establishes commensal microorganisms which develop into the gut microbiome [18]. To understand how Lm is able to access and cross the intestinal epithelium to establish hematogenous infection, it is crucial to understand microbial interactions within the maternal gut environment and how this microbiome influences Lm pathogenicity. While many studies exist on understanding the intracellular phase of Lm infection, very little is known about the behavior of Lm within the gastrointestinal tract. Few studies have characterized microbial dysbiosis during listeriosis [19]. It is possible that the maternal gut microenvironment may play a role in dispersion of Lm outside of the intestinal tract, with commensal microbes influencing Lm survival and invasion of epithelial tissues. We hypothesized that pregnancy is associated with the hematogenous spread and severity of Lm infection through dysbiosis of the homeostatic gut microbiome that does not occur in nonpregnant hosts. To test this hypothesis, we characterized the gut microbiota of both pregnant and nonpregnant NHP following challenge with Lm using 16S rRNA sequencing.

## Ethics statement

The rhesus macaques used in this study were cared for by the staff at the WNPRC in accordance with the regulations and guidelines outlined in the Animal Welfare Act and the Guide for the Care and Use of Laboratory Animals and the recommendations of the Weatherall report [20]. Per WNPRC standard operating procedures for animals assigned to protocols involving the experimental inoculation of an infectious pathogen, environmental enhancement included constant visual, auditory, and olfactory contact with conspecifics, the provision of feeding devices which inspire foraging behavior, the provision and rotation of novel manipulanda (e.g., Kong toys, nylabones, etc.), and enclosure furniture (i.e., perches, shelves). Per Animal Welfare Regulations (Title 9, Chapter 1, Subchapter A, Parts 1–4, Section 3.80 Primary enclosures) animals were housed in a nonhuman primate Group 3 enclosure with at least 4.3 square feet of floor space and at least 30 inches of height. This study was approved by the University of Wisconsin-Madison Graduate School Institutional Animal Care and Use Committee (animal protocol number 005061).

All animals were housed in enclosures with at least 4.3, 6.0, or 8.0 sq. ft. of floor space, measuring 30, 32, or 36 inches high, and containing a tubular PVC or stainless-steel perch. Each individual enclosure was equipped with a horizontal or vertical sliding door, an automatic water lixit, and a stainless-steel feed hopper. All animals were fed using a nutritional plan based on recommendations published by the National Research Council. Twice daily, macaques were fed a fixed formula, extruded dry diet (2050 Teklad Global 20% Protein Primate Diet) with adequate carbohydrate, energy, fat, fiber (10%), mineral, protein, and vitamin content. Dry diets were supplemented with fruits, vegetables, and other edible foodstuffs (e.g., nuts, cereals, seed mixtures, yogurt, peanut butter, popcorn, marshmallows, etc.) to provide variety to the diet and to inspire species-specific behaviors such as foraging. To further promote psychological well-being, animals were provided with food enrichment, human-to monkey interaction, structural enrichment, and manipulanda. Environmental enrichment objects were selected to minimize chances of pathogen transmission from one animal to another and from animals to care staff. While on study, all animals were evaluated by trained animal care staff at least twice daily for signs of pain, distress, and illness by observing appetite, stool quality, activity level, physical condition. Animals exhibiting abnormal presentation for any of these clinical parameters were provided appropriate care by attending veterinarians. Prior to all minor/brief experimental procedures, animals were sedated using ketamine anesthesia, which was reversed at the conclusion of a procedure using atipamizole. Animals undergoing surgical delivery of fetuses were pre-medicated with ketamine and general anesthesia was maintained during the course of the procedure with isoflurane gas using an endotracheal tube. Animals were monitored regularly until fully recovered from anesthesia. Delivered fetuses were anesthetized with ketamine, and then euthanized by an intramuscular or intraperitoneal overdose injection of sodium pentobarbital.

## Methods

### Care and Use of Macaques

Female cynomolgus macaques were housed and cared for by Wisconsin National Primate Research Center (WNPRC) staff in accordance with the regulations and guidelines outlined in the Animal Welfare Act and the Guide for the Care and Use of Laboratory Animals. All animals were systematically monitored twice daily by WNPRC staff and veterinarians, and additionally as needed. All observations were entered into the colony electronic health records. Menstrual cycle monitoring was performed through daily monitoring and vaginal swabbing by WNPRC animal care. Blood samples were collected using a needle and syringe or vacutainer system from the femoral or saphenous vein. This study was approved by the University of Wisconsin-Madison College of Letters and Sciences and the Vice Chancellor Office for Research and Graduate Education Institutional Animal Care and Use Committee (IACUC).

### Cesarean section and tissue collection (fetectomy)

All fetal and maternal tissues were surgically removed at laparotomy. These were survival surgeries for the dams. The entire conceptus within the gestational sac (fetus, placenta, fetal membranes, umbilical cord, and amniotic fluid) was collected and submitted for necropsy. The fetus was euthanized with an overdose of sodium pentobarbital (50 mg/kg). Tissues were carefully dissected using sterile instruments that were changed between each organ and tissue type to minimize possible cross contamination. Each organ/tissue was evaluated grossly in situ, removed with sterile instruments, placed in a sterile culture dish, and sectioned for histology, bacterial burden assay, or banked for future assays. Biopsies of the placenta, decidua, maternal liver, spleen, and a mesenteric lymph node were collected aseptically during surgery into sterile petri dishes, weighed, and further processed for bacterial burden and histology. Maternal decidua was dissected from the maternal surface of the placenta.

### Necropsy and tissue collection

For terminal studies, animals were euthanized with an overdose of sodium pentobarbital (50 mg/kg). At 14 days post infection (dpi), non-pregnant subjects were sedated, euthanized, and sterile instruments were used for the dissection and collection of colon, cecum, jejunum, and uterine epithelium tissues during the gross post-mortem examination. Each tissue was evaluated grossly in situ, removed with sterile instruments, placed in a sterile culture dish, and sectioned for histology, bacterial burden assay, or banked for future assays.

### Subjects

Thirty-two adult female cynomolgus macaques (*Macaca fascicularis*) were used in this study (Supplementary Table 1). Four of the eight pregnant cohort of Lm-inoculated animals have previously been described [1]. Animals were housed in group 3 or group 4 enclosures in accordance with the Animal Welfare Act and its regulations and the Guide for the Care and Use of Laboratory Animals. All animals were monitored twice daily by an animal researcher or veterinary technician for evidence of disease or injury. Body weight was monitored to ensure that all animals remained in properly sized cages. Animals were fed commercial nonhuman primate chow (2050 Teklad Global 20% Protein Primate Diet, Harlan Laboratories, Madison, WI) twice daily, supplemented with fruits or vegetables and a variety of environmental enrichment. All animals used were actively mensing and had not entered menopause. Menstrual cycle assessment was performed through daily monitoring and vaginal swabbing by WNPRC animal care personnel. The age of subjects ranged from 5-13 years.

### *Listeria* Inoculation

The methodology and outcomes of Lm inoculation during pregnancy for subjects included in this study have been previously published[2]. Some of these animals were subsequently included as nonpregnant subjects in the current study. No antibiotics were administered to any animals during the course of pregnant or nonpregnant inoculation.

*Listeria monocytogenes* (Lm; Lm2203 [21]) was cultured at 37 °C in Tryptic Soy Broth (Becton Dickinson, Sparks, MD). Each inoculum containing 1 x 10^8^ colony forming units (CFU)/ml of Lm at log-phase growth was diluted in 10mL of whipping cream and delivered via oral gavage through a soft intragastric feeding tube under sedation (n=16), as previously described [1, 2, 22]. Control inoculations (mock) consisted of 10mL of whipping cream alone with no Lm (n=14). These 30 subjects are organized into four cohorts (Supplementary Table 1), with Cohort 1 including non-pregnant controls, Cohort 2 including non-pregnant Lm-exposed females, Cohort 3 including pregnant control dams, and Cohort 4 including pregnant Lm-exposed dams.

To confirm the dose of Lm given to each subjects, 500 μL of the 10mL whipping cream inoculum was serially diluted in phosphate-buffered saline (PBS; Catalog #P5368, Sigma-Aldrich, St. Louis, MO), plated on Trypticase soy agar with 5% sheep blood (Becton Dickinson, Sparks, MD), and quantified after overnight incubation at 37^⍰^C.

### Fecal Shedding

Lm fecal shedding and bacteremia were evaluated during the 14-day period following Lm inoculation (Supplemental Fig. 1), as previously described [1, 2]. Fecal samples were collected from cage pans daily, starting on the day of inoculation prior to the first dose of inoculum being given and ending on the day of tissue collection. Samples were collected from cage pans. Serial fecal dilutions in PBS were plated on Modified Oxford Medium (Fischer Scientific Hampton, NH) and incubated at 37^⍰^C for 48 hours to identify and quantify Lm. The number of colonies was quantified at both 24 hours and 48 hours after plating.

### Diarrhea Observation and Scoring

Fecal material was assessed for diarrhea and numerical values were assigned as follows: 0 = no abnormal observations, 1= soft feces, 2 = diarrhea, and 3= wet diarrhea.

### Bacteremia

Whole peripheral blood samples were collected every 2-3 days and processed by the Clinical Pathology Laboratory at the School of Veterinary Medicine at the University of Wisconsin-Madison[1, 2]. BD Bactec Peds Plus/F blood culture bottles (Becton Dickinson Diagnostic Systems, Sparks, MD) were aseptically inoculated with 3 mL whole blood per bottle. The samples were then incubated at 35°C in a BD Bactec 9050 blood culture system (Becton Dickinson Diagnostic Systems, Sparks, MD) until a positive signal was observed or for a maximum of 5 days. Recovered isolates were identified by matrix-assisted laser desorption-ionization time-of-flight (MALDI-TOF) mass spectrometry (Bruker Daltonics, Billerica, MA). Sample extraction and strain identification was performed following manufacturer’s instruction. A score of >2 indicated genus and probable species identification.

### Tissue Collection and Processing

Tissues from Cohorts 1 and 2 nonpregnant monkeys were collected at 14 dpi. The monkeys were anesthetized with ketamine hydrochloride (10-15 mg/kg, iv) and euthanized with an overdose of pentobarbital sodium (a minimum of 25 mg/kg, iv). The uterus and selected segments of the GI tract were removed, and the endometrium was scraped from the myometrium for analysis. The colon, cecum, jejunum, and endometrium were collected in addition to liver, spleen, and lymph nodes (Supplemental Table 2). Segments of collected tissues were fixed and embedded for histology (Supplemental Fig 3) or homogenized for bacteriological analysis on blood agar plates as previously described [23].

Tissue collection from Cohorts 3 and 4 p(regnant monkeys) was described previously [1, 2]. Briefly, following inoculation, if fetal demise was indicated by absence of heartbeat, fetal and maternal tissues were promptly collected at laparotomy. The placenta, decidua, and fetal tissues were collected in addition to maternal liver, spleen, and lymph nodes (Supplemental Table 2).

### Histology

Tissues collected for histology were fixed in 4% PFA overnight followed by 70% ethanol overnight, and then processed and embedded in paraffin. 5µm sections were stained with H&E and assessed by veterinary pathologists blinded to treatment groups. Tissues were evaluated for the presence or absence of pathologic changes, normal anatomic variations, and inflammation. Organs considered to have no significant pathologic or inflammatory changes and were scored as 0. Severity (Supplemental Table 3) was determined by the extent and distribution of inflammation, vascular change (infarction, thrombosis, pregnancy-associated vascular remodeling and/or the lack thereof), and non-vascular necrosis across the tissue section or organ (multiple slides were necessary to evaluate the placenta). Scores were averaged and compared between treatment groups as previously reported [24].

### DNA extraction, PCR, and sequencing

Fecal samples were analyzed at 4 timepoints from all but one individual animal in the 4 cohorts (0, 3-5, 7-10, and 14 dpi); two samples were lost for one animal in the pregnant control cohort (ID 24A & 24B). The DNA extraction methods utilized in this analysis were previously described in detail [25]. Bacterial DNA was isolated using a Qiamp PowerFecal DNA Isolation Kit (Qiagen, Hilden, Germany). A negative control was inserted periodically in the workflow after blocks of 16 samples to test for methodological contamination during processing. All negative controls yielded an undetectable amount of DNA. DNA was quantified on a Qubit 2.0 Fluorometer (Thermo Fisher Scientific, Waltham, MA, USA) using Qubit fluorometric quantitation reagents (Thermo Fisher Scientific, Waltham, MA, USA).

The fourth hypervariable (V4) region of the bacterial 16S rRNA gene was amplified using the one-step polymerase chain reaction (PCR) approach with barcoded V4 primers (F- GTGCCAGCMGCCGCGGTAA; R- GGACTACHVGGGTWTCTAAT). Each primer pair was barcoded with individual custom indices to facilitate demultiplexing, as previously described [26]. Each PCR reaction consisted of 12.5 μl KAPA 2x HiFi Master Mix (KAPA Biosystems, Wilmington, MA, USA), 0.5 μl of 10 μM forward primer, 0.5 μl of 10 μM reverse primer and up to 11.5 μl of 10ng/μl DNA to a total volume of 25 μl with nuclease-free water (IDT, Coralville, Iowa, USA). Amplification conditions on a C1000 Touch™ thermal cycler (Bio-Rad Laboratories, Hercules, CA, USA) were 95°C for 3 min, 35 cycles of 95° for 30 s, 55°C for 30 s, and 72°C for 30 s, followed by a final extension at 72°C for 5 min. The PCR products were purified by running on a 1% low-melt agarose gel (National Diagnostics, Atlanta, GA) stained with SYBR Safe DNA Gel Stain (Invitrogen, Waltham, CA) to isolate amplicons of the expected size (∼380 bp). DNA bands at ∼380 bp were excised and purified utilizing the Zymo Gel DNA Recovery Kit (Zymo Research, Irvine, CA, United States). Purified PCR products were equimolar pooled for a final library concentration of 11 pmol/l. Sequencing was performed on an Illumina MiSeq (Illumina, San Diego, CA, USA) with 10% PhiX control using a 500-cycle v2 (2×250 paired-end) sequencing kit and custom sequencing primers [26].

### 16S rRNA sequencing data processing

Raw sequencing data were processed using mothur [27] (version 1.43.0) and Qiime 2 [28] . Contigs (overlapping sequences) were aligned using the SILVA database (v132) [29] and low-quality reads and chimeras were detected by UCHIME and removed. Sequences were assigned to operational taxonomic units (OTUs) with a threshold of 97% similarity using the SILVA database. OTUs with less than 0.01% overall abundance within the dataset were considered rare and were removed from the dataset. After rare OTUs were filtered, each sample each sample was subsampled to 3,200 reads to normalize against the sample with the lowest number of sequences.

### Statistical analysis

Normalized OTU counts were used to determine diversity metrics. Diversity metrics were calculated for all samples using Qiime and RStudio (v2023.5 and v2023.03 respectively) [30]. All alpha- (within sample diversity) and beta-diversity (between sample comparisons) metrics and relative abundance measures were calculated using the phyloseq package in R [31].

To assess the stability of alpha-diversity measures by infection status, we compared the mean (by cohort) of observed OTUs using Shannon’s Diversity Index. We additionally assessed the change in microbial composition between cohorts, examining both pregnancy/nonpregnancy and infection status separately. To meet the objective of determining changes in the microbiota following experimental challenge, microbiota composition, alpha diversity metrics, the Bray-Curtis [32] and weighted Unifrac [33] beta diversity metrics, and the relative abundance of dominant genera were compared between cohorts, reproductive state, and APO. Common dominant genera within Cohort 4 were evaluated for effects of bacteremia, tissue infection with Lm, and APO on the RA using the generalized linear mixed models described above, accounting for calf as a random effect. Alpha was set at 0.05 for all statistical analyses.

Differences in the Bray-Curtis and weighted Unifrac index between groups were used to create a sample-wise distance matrix that was visualized using multidimensional scaling (MDS). Non-metric MDS (NMDS) was used if the base stress of creating the ordination plots was non-linear; metric MDS was utilized if the base stress followed linear regression. Equality of beta dispersion between groups was assessed using the betadisper function of the vegan package. Bray-Curtis and weighted Unifrac was compared between treatment groups (i.e., Cohorts), Reproductive State, and APOs using a permutational multivariate analysis of variance (PERMANOVA) as implemented in the vegan package.

Statistical analysis for observational data including diarrhea, weight change, and tissue pathology were performed using GraphPad Prism version 9.0.0 for Mac GraphPad Software, San Diego, California USA (http://www.graphpad.com).

## Results

We utilized a non-human primate model of listeriosis during early gestation and normal menstrual cycles. The well-being of experimental subjects was monitored by evaluation of weight during the period of study. One-way ANOVA followed by Tukey’s multiple comparisons test revealed no significant difference in weight change (between timepoint A & D) among the experimental cohorts (Fig 1a). Subjects were also monitored closely for signs of gastrointestinal upset. The incidence and severity of diarrhea was monitored in all groups (Fig 1b). While there was occasional diarrhea in some of the subjects, there were no significant differences between groups with comparison by exposure to Lm or during pregnancy (Fig 1d). Along with daily fecal sample collections, feces type and severity were monitored closely (Fig 1b). Scoring was conducted as mentioned above (See Methods). Statistical analysis of collected data using three-way ANOVA followed by Tukey’s multiple comparisons test revealed no significant associations of diarrhea severity with treatment group, reproductive state, Lm exposure, or subject (Supplemental Table 4).

**Figure 1a.**
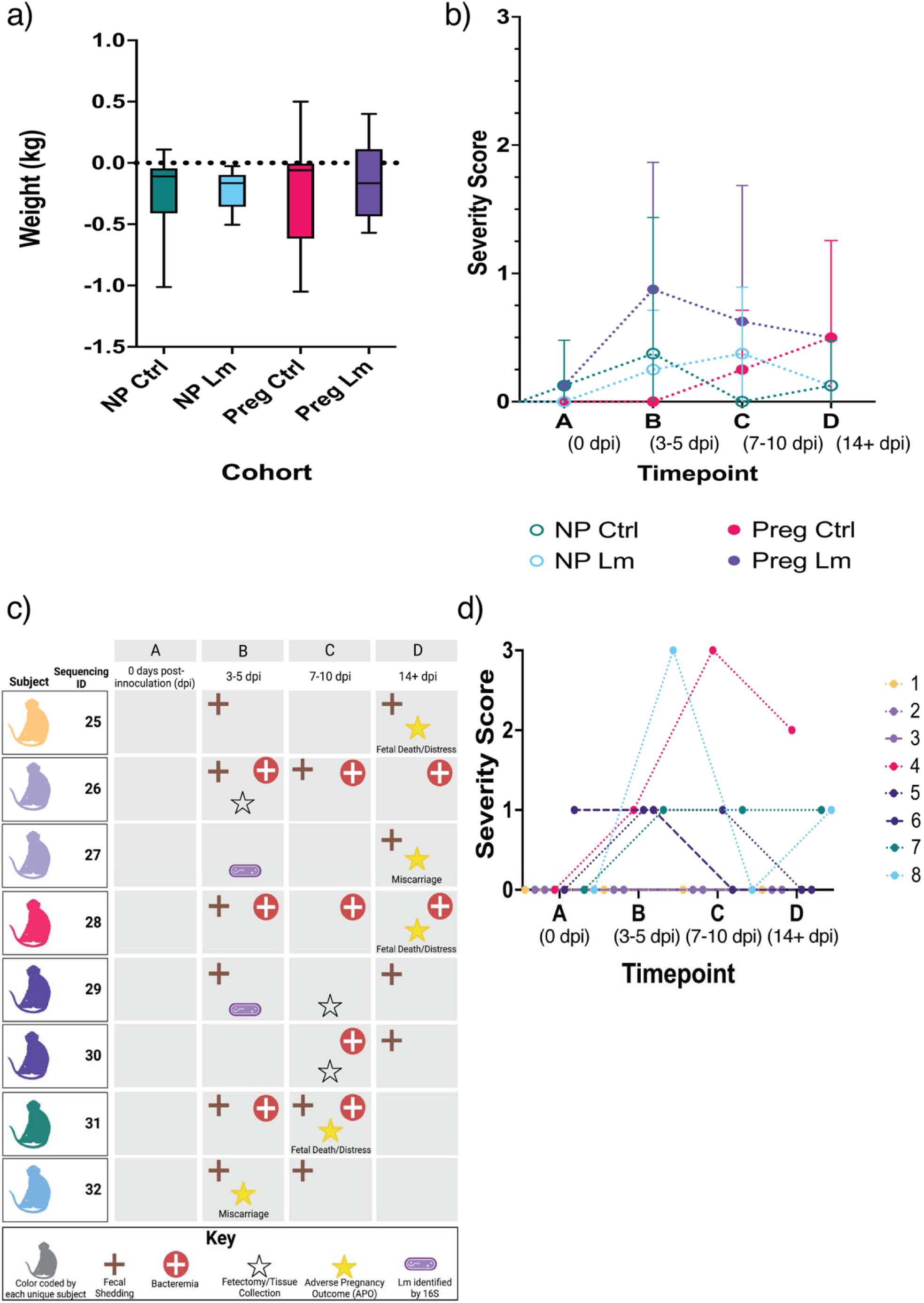
A violin plot depicting the average change in weight between timepoint A and D. Each cohort is color coded; these colors are carried through subsequent figures. The mean weight change is indicated by the horizontal line in each plot and standard error of the mean (SEM) denoted by bars. **Figure 1b.** A graph with the average diarrhea severity score of each cohort on the y-axis and the timepoint (A-D) during the experiment on the x-axis. Average values are depicted by circles and SEM denoted by bars. Each cohort is color coded. A = pre-inoculation, B = 3-5 days following inoculation, C = 7-10 following inoculation, and D = tissue collection and conclusion of the experiment. **Figure 1c.** Summary table of fecal shedding of Lm, bacteremia, and occurrence pregnancy outcomes of each subject in the pregnant Lm cohort. Each subject is color coded. Some subjects were utilized in the experimental protocol twice and share a color code. The Key at the bottom explains each symbol. The table depicts fecal and bacterial shedding of Lm throughout timepoints A-D, from left to right. The outcome of each pregnancy is denoted by a hollow star indicating tissue collection and a filled star indicating APO. Furthermore, those subject with Lm identified by sequencing are indicated by a bacterial rod. **Figure 1d**. Diarrhea severity score of each subject within Cohort 4 plotted against the experimental timepoint (A-D). Each subject is color coded. The dashed lines indicate the same subject utilized a second time within the experimental protocol A = pre-inoculation, B = 3-5 days following inoculation, C = 7-10 following inoculation, and D = tissue collection and conclusion of the experiment.

To characterize the progress of listeriosis and dissemination of bacteria, we monitored fecal shedding, bacteremia, bacterial burden within tissues, and pregnancy outcomes. The non-pregnant cohort displayed no fecal shedding, dissemination, or pathology. As only the pregnant Lm-exposed cohort had observable of listeriosis, we summarized and organized their data in Fig 1c, which lists the 8 experimental subjects in the Pregnant Lm cohort on the left. Within this cohort, all subjects shed Lm in the feces at some time during the 2 weeks following inoculation. Four of the eight subjects had bacteremia, and five of the eight subjects had fetal demise or miscarriage (Fig 1c). While fecal Lm was identified by culture-based methods in all of Cohort 4, Lm was identified by sequencing-based methods in the same individual who was utilized twice in this protocol (subjects 27 & 28), and both samples were during 3-5 dpi (Supplemental Fig 4).

Although all animals in Cohort 4 had identifiable fecal carriage of Lm, using the diarrhea scoring mentioned previously (Fig 1d) revealed no association of diarrhea severity with subject or timepoint (A-D) (Sup Table 4).

To assess the impact of Lm dissemination, we statistically evaluated pathology in Lm target tissues (liver, spleen, lymph nodes) from all experimental cohorts, as well as colon, cecum, jejunum, and endometrial lining from the non-pregnant cohorts and the decidua, placenta, and fetal tissues for the pregnant cohorts (Supplemental Fig 3). Pathology severity was assessed by ACVP board-certified WNPRC Veterinary Pathologists and assigned a score based on our previously documented rubric (Sup Table 3)[2]. We evaluated the data using 3-way ANOVA statistical analysis to assess any significant associations among exposure, reproductive state, and tissue pathology. As expected, based on previously published data, there was a significant increase in the pathology scores in the decidua, placenta and fetal tissues for Lm infected versus uninfected animals [1]. In the nonpregnant cohort, Lm had no effect on the endometrium or gastrointestinal tissues. Although there were no significant differences in histopathology in the spleen, there were significantly lower liver and lymph node pathology scores in non-pregnant Lm-exposed subjects, compared to their pregnant counterparts. The reasons for these isolated differences are not known at this time.

From the collected feces, a total of 128 samples were sequenced. Bacterial amplicon sequencing of the 16S rRNA gene generated a total of 3,656,601 raw sequences with an average of 28,567 ± 3,505 sequences per sample (mean ± SE; range 7,757–13,859). Sequence clean-up in Qiime2 resulted in a total of 3,651,002 sequences for an average of 32,589 ± 3,312 sequences per sample (range 19,172–25,796). After normalization, 112 samples remained consisting of 30 non-pregnant controls, 29 non-pregnant Lm-exposed, 23 pregnant controls, and 29 pregnant Lm-exposed.

Figure 2 presents the 25 most abundant OTUs to the highest level of taxonomic identification. There was considerable inter- and intra-animal individuality. The data indicate a high abundance of *Prevotella* spp. and *Treponema* spp. which are known to be predominant in NHPs, compared to humans[34]. The other abundant OTUs are similar to those found in the human gut microbiome [35], indicating that this a highly translatable model.

**Figure 2.**
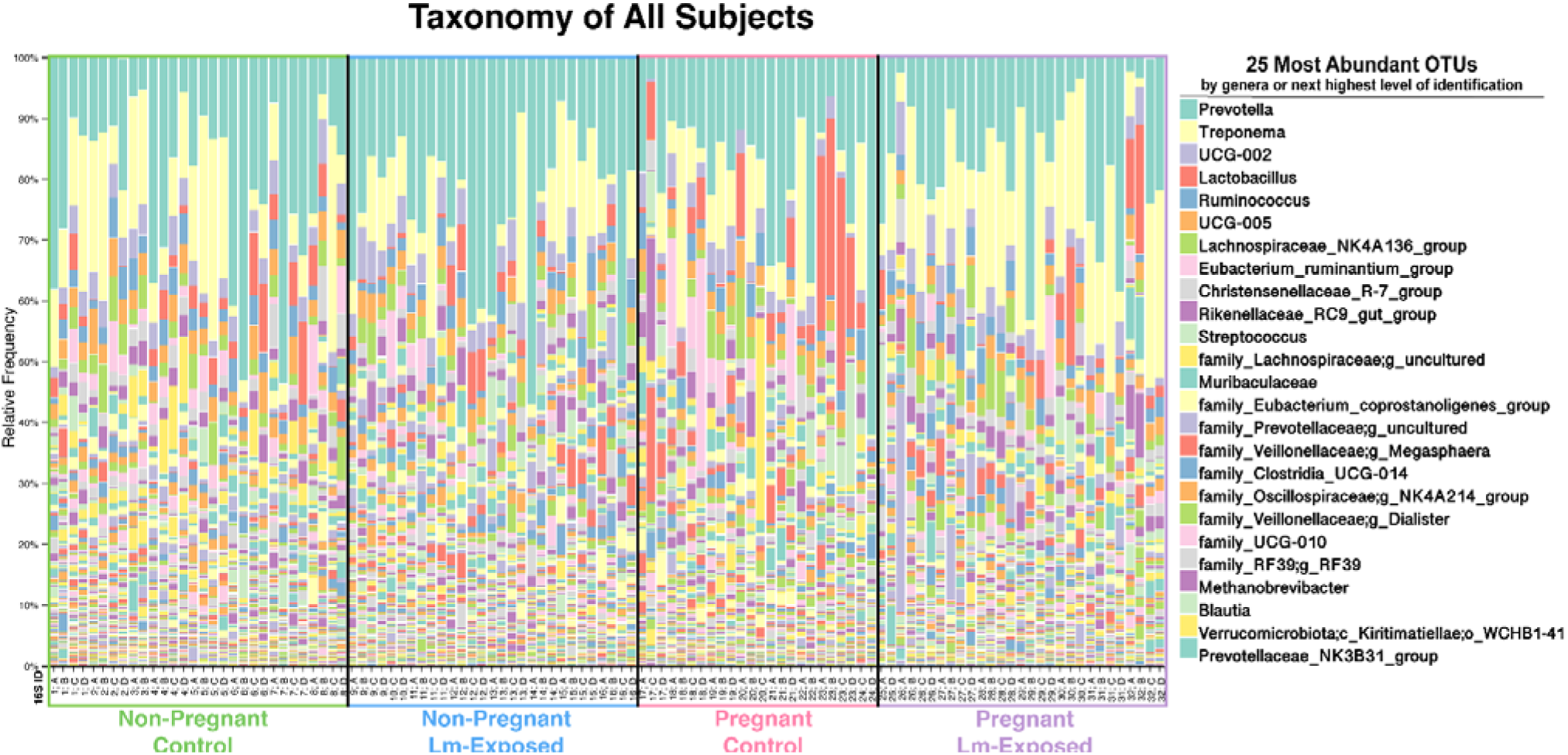
Taxonomy Bar plot of 25 most abundant OTUs in fecal samples from experimental cohorts. Each cohort is indicated along the x-axis, and the samples from individual animals as described in Fig. 1 are presented chronologically (e.g., samples 1A, 1B, 1C and 1D are the first presented, followed by 2A, 2B etc.). The top 25 most abundant OTUs are listed at the right in order of highest abundance and classified by the highest level of identification. Each cohort is separated by black bars.

We evaluated the community richness of our maternal intestinal environments by determining alpha-diversity. Alpha-diversity is a measure of the relative abundances of the different species that make up the richness of a sample and is represented by assigning a value (H-value) for the species in a particular ecosystem. There are several metrics by which alpha-diversity can be measured. We utilized the Shannon Diversity Index to estimate species richness and evenness or the average diversity of a species within a sample on a local scale. Figure 3 illustrates the Shannon Entropy Index (alpha diversity) of the fecal microbiota in the four experimental cohorts. The results demonstrate that the non-pregnant control and non-pregnant Lm-exposed groups were significantly different from the Pregnant Lm-exposed group, but not significantly different from the pregnant control group, suggesting an interaction between the pregnant state and susceptibility to gut dysbiosis during listeriosis (Fig 3a). The data also indicate that Lm by itself does not impact the gut microbiome in the nonpregnant state. Rather, the combined nature of pregnancy and exposure to Lm was associated with significant loss in community richness and diversity (Fig 3b). Furthermore, when examining the Lm-exposed cohort only, there is a significant loss in diversity in the pregnant Lm-exposed vs non-pregnant Lm-exposed cohorts. This confirms the importance of pregnancy in the susceptibility of the pregnant state to dysbiosis with Lm exposure. When examining only the control cohorts, there was no significant impact of pregnancy on alpha diversity (Supplemental Fig 5), underscoring the interaction between reproductive state and Lm exposure.

**Figure 3a.**
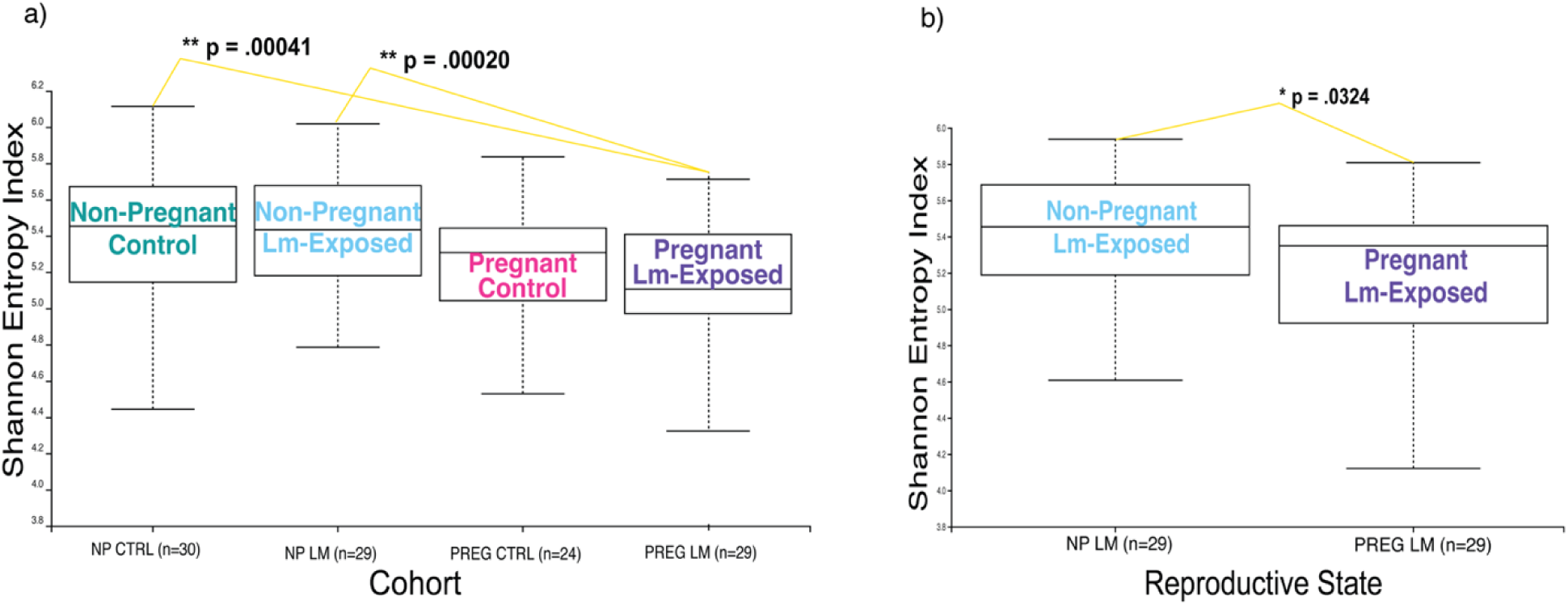
Box plot of the alpha-diversity measure by Shannon Entropy Index of the fecal microbiota in the four experimental cohorts. The average H-value is presented with bars indicating SEM. Sample size is listed on the x-axis. Significance is denoted by asterisks and the significantly different cohorts are connected by yellow lines. * P ≤ 0.05, ** P ≤ 0.01. **Figure 3b** Box plot of the alpha-diversity measure by Shannon Entropy Index of the fecal microbiota in the Lm-exposed experimental cohorts only. The average H-value is presented with bars indicating SEM. Sample size is listed on the x-axis. Significance is denoted by asterisks and the significantly different cohorts are connected by yellow lines. * P ≤ 0.05, ** P ≤ 0.01.

We also evaluated whether there was a discernable time-related impact of exposure to Lm on gut microbial richness. Although there was no significant dysbiosis in the pregnant Lm-exposed cohort following introduction of Lm (Sup. Fig 6), there was a statistically significant difference between timepoints C (7-10 days post-inoculation) and D (14+ days post-inoculation) within the nonpregnant Lm-exposed cohort (Sup. Fig. 5). Perhaps the increase in alpha diversity at 7-10 dpi represents a restoration of diversity following resolution of Lm.

**Figure 4.**
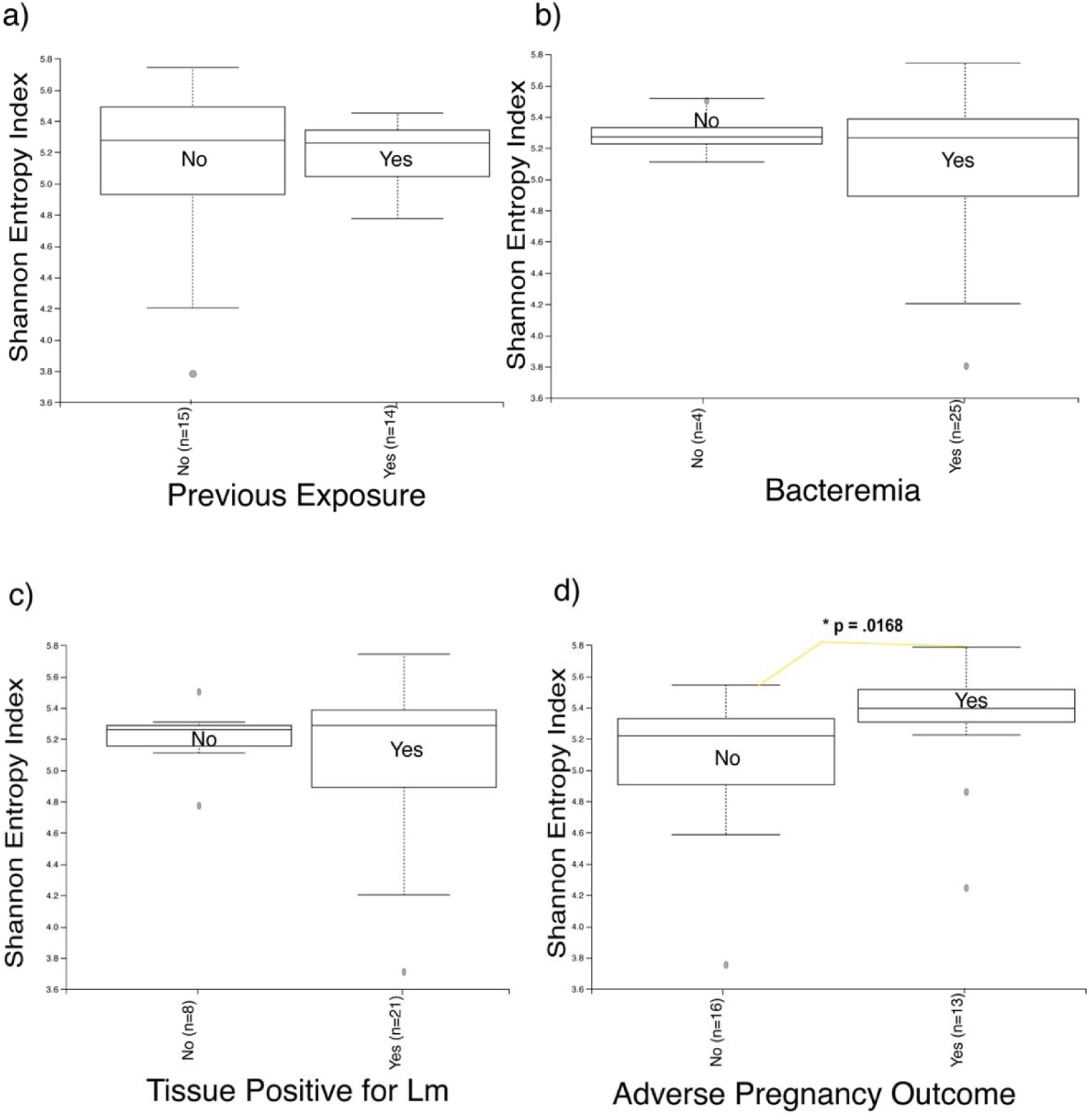
Alpha Diversity measured by Shannon Entropy Index of the Pregnant Lm-Exposed cohort evaluated with regard to previous exposure(4a), bacteremia(4b), tissue bacterial burden(4c), and adverse pregnancy outcome(4d). Fecal shedding of Lm was not analyzed as all subjects were positive for shedding. The average H-value is presented as box plots with bars indicating SEM. “No” indicates negative and “Yes” indicates positive for Previous Exposure, Bacteremia, Tissue positive for Lm, or occurrence of APO; the respective n is listed on the x-axis. Outliers are marked by a filled circle. Significance is denoted by asterisks and the significantly different cohorts are connected by yellow lines. * P ≤ 0.05, ** P ≤ 0.01.

**Figure 5a.**
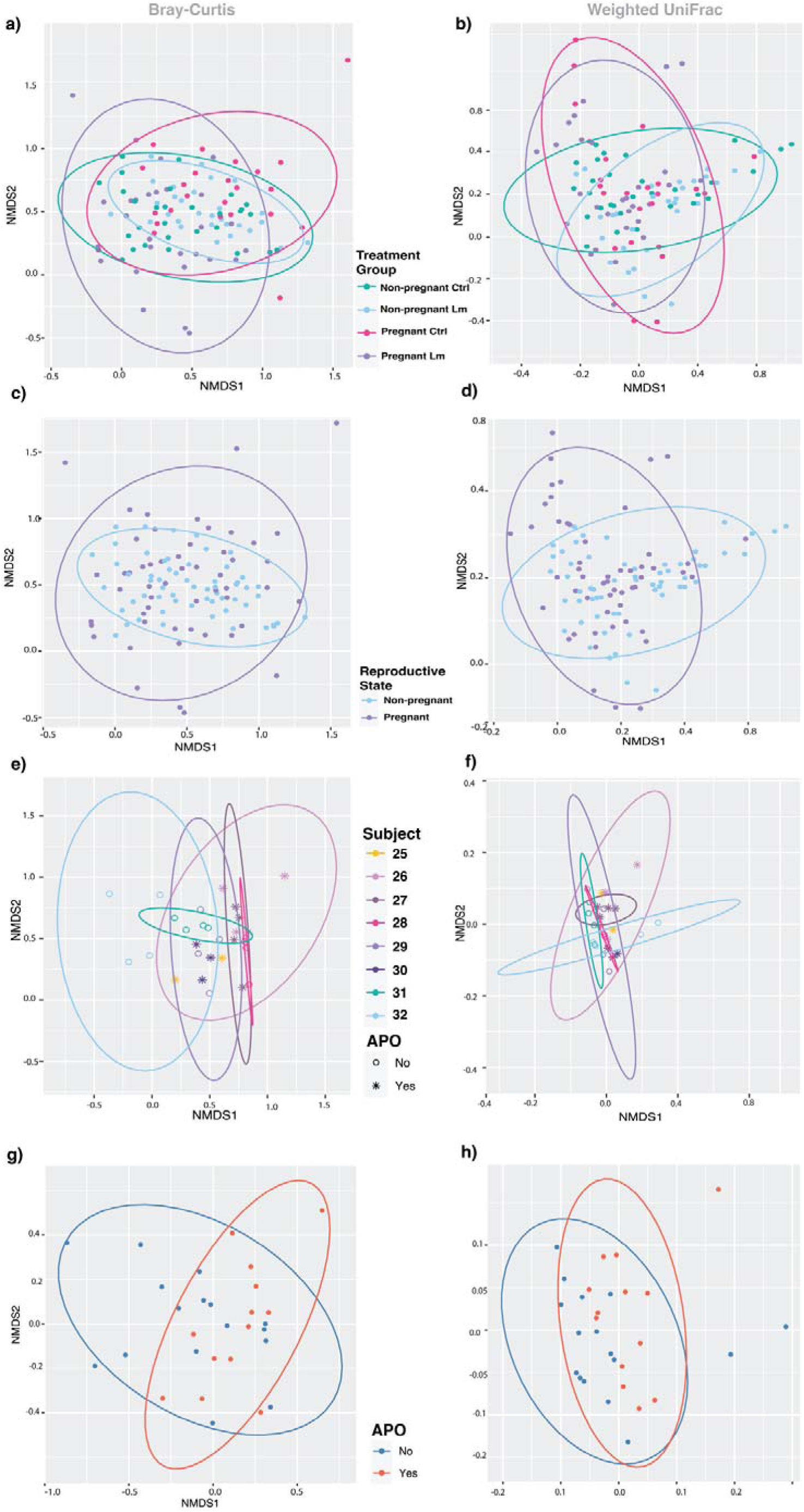
Beta Diversity by Treatment Group, depicted via principal coordinate analysis (PCA) calculated using Bray-Curtis dissimilarity matrix at Genus level abundances. Ellipses are color coded by cohort and depict 95% confidence grouping. **Figure 5b** Beta Diversity by Treatment Group, depicted via principal coordinate analysis (PCA) calculated using weighted UniFrac analysis at Genus level abundances. Ellipses are color coded by cohort and depict 95% confidence grouping. **Figure 5c** Beta Diversity by Reproductive State, depicted via principal coordinate analysis (PCA) and calculated using Bray-Curtis dissimilarity matrix at Genus level abundances. Ellipses are color coded by cohort and depict 95% confidence grouping. **Figure 5d** Beta Diversity by Reproductive State, depicted via principal coordinate analysis (PCA) calculated using weighted UniFrac analysis at Genus level abundances. Ellipses are color coded by cohort and depict 95% confidence grouping. **Figure 5e** Beta Diversity by Subject of the pregnant Lm-exposed cohort depicted via principal coordinate analysis (PCA) and calculated using Bray-Curtis dissimilarity matrix at Genus level abundances. Occurrence of APO is noted by an asterisk shape. Ellipses are color coded by cohort and depict 95% confidence grouping. **Figure 5f** Beta Diversity by Subject of the pregnant Lm-exposed cohort depicted via principal coordinate analysis (PCA) and calculated using weighted UniFrac dissimilarity matrix at Genus level abundances. Occurrence of APO is noted by an asterisk shape. Ellipses are color coded by cohort and depict 95% confidence grouping. **Figure 5g** Beta Diversity by APO, depicted via principal coordinate analysis (PCA) calculated using Bray-Curtis analysis at Genus level abundances. Occurrence of APO is noted by an asterisk shape. Ellipses are color coded by cohort and depict 95% confidence grouping. **Figure 5h** Beta Diversity by APO, depicted via principal coordinate analysis (PCA) calculated using weighted UniFrac analysis at Genus level abundances. Occurrence of APO is noted by an asterisk shape. Ellipses are color coded by cohort and depict 95% confidence grouping.

**Figure 6a.**
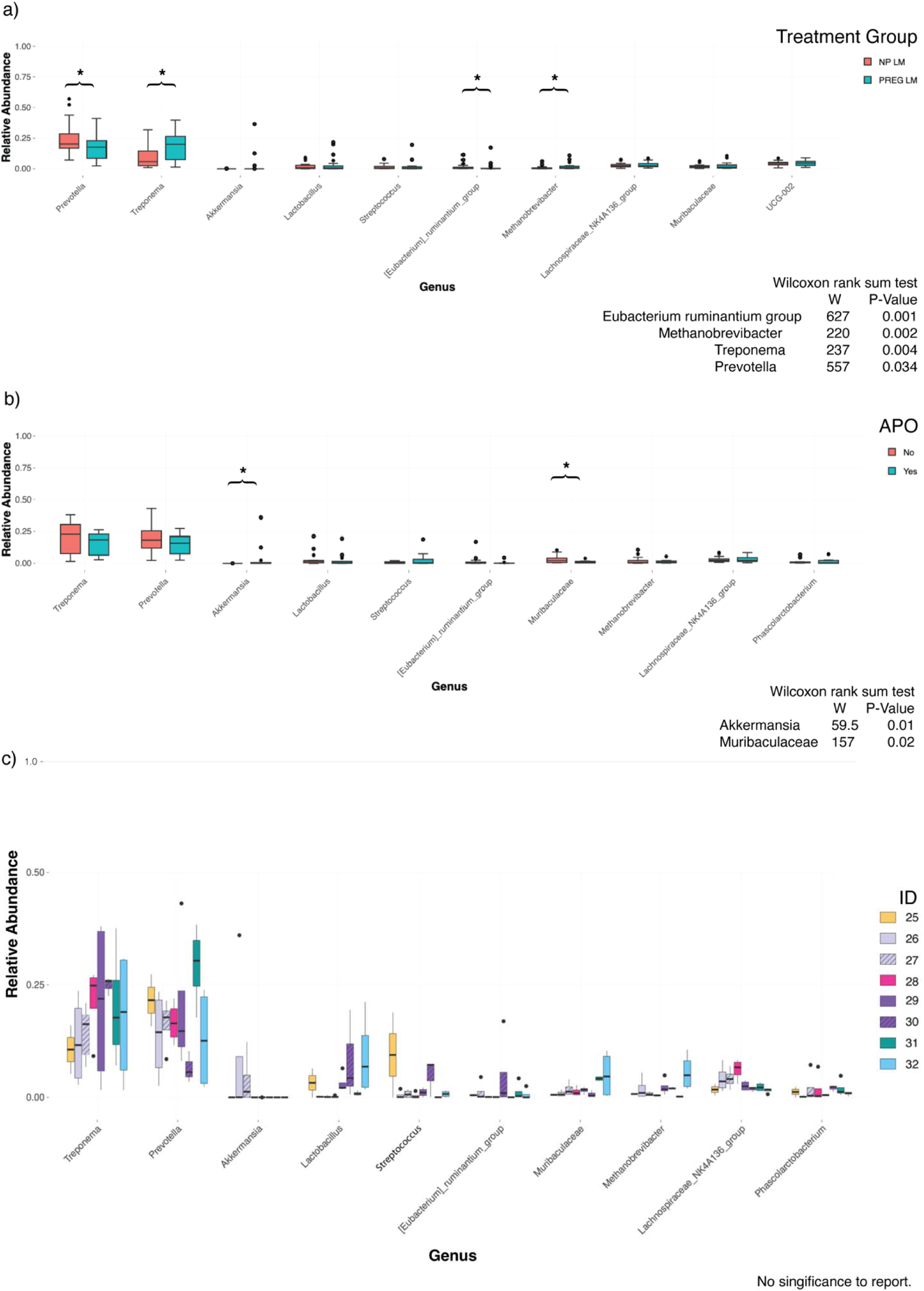
The box plot illustrates the variation of abundance reads of the ten most abundant taxa across all timepoints (A-D) of the Lm-exposed Cohort 2 & 4. Reproductive state is color coded. OTUs are listed left to right, in order of decreasing abundance. The average abundance of each OTU is indicated by a black line in the middle of the bars and SEM denoted by lines. Outliers are represented by black dots. Significance is denoted by asterisks. * P ≤ 0.05, ** P ≤ 0.01. **Figure 6b** The box plot illustrates the variation of abundance reads of the most abundant taxa across all timepoints (A-D) of the pregnant Lm-exposed Cohort 4. The occurrence of APOs is color coded. OTUs are listed left to right, in order of decreasing abundance. The average abundance of each OTU is indicated by a black line in the middle of the bars and SEM denoted by lines. Outliers are represented by black circles. Significance is denoted by asterisks. * P ≤ 0.05, ** P ≤ 0.01. **Figure 6c** The box plot illustrates the variation of abundance reads of the most abundant taxa across all timepoints (A-D) of each subject in the pregnant Lm-exposed Cohort 4. OTUs are listed left to right, in order of decreasing abundance. Subjects are color coded. Those subjects which were used twice in the experiment are indicated with black diagonal stripes within the bars. The average abundance of each OTU is marked by a black line in the middle of the bars and SEM denoted by lines. Outliers are represented by black dots.

Because the pregnant Lm-exposed cohort 4 was the only cohort to exhibit signs of listeriosis, we evaluated alpha diversity of Cohort 4 in the in regard to pregnancy outcomes, bacteremia, tissue bacterial burden, and previous exposure to Lm. There were no significant changes in alpha diversity associated with previous exposure, bacteremia, or bacterial burden (Fig 4a-c). However, there was a significant increase in alpha diversity in those individuals who had APOs (miscarriage, intrauterine fetal demise) compared to subjects that did not (Fig 4d).

To determine whether our treatment groups were heterogeneous, we next examined beta-diversity which measures the distance or dissimilarity between each sample pair. Similar to alpha-diversity, there are several metrics for calculating beta-diversity. For our analysis, the Bray–Curtis dissimilarity and weighted UniFrac were employed, as they quantify the compositional dissimilarity between sites based upon relative abundance. UniFrac differs from Bray-Curtis in that it incorporates phylogenetic distances and allows for the option to consider the relative abundance of taxa shared between samples (weighted vs. un-weighted). These statistical analyses are organized in Supplementary Table 4. We then visualized both Bray-Curtis and weighted UniFrac metrics via principal coordinate analysis (PCA) (Fig 5).

We began by examining all cohorts. Unsupervised clustering and analysis revealed that the data cluster by treatment group (betadisper adj P = 0.001) and that microbial composition was significantly dissimilar between cohorts (P = 0.001) (Figure 5a). Weighted clustering and analysis confirmed clustering (betadisper adj P = 0.005), but indicated that the microbial composition between treatment groups was similar when accounting for abundances (Figure 5b).

When evaluating the Lm-exposed cohorts, unsupervised clustering and analysis supported clustering by reproductive state (betadisper adj P = 0.005) and microbial composition dissimilarity between the pregnant and non-pregnant state (P = 0.001) (Fig 5c). Weighted clustering and analysis by reproductive state confirmed clustering (betadisper adj P = 0.007) but indicated that pregnancy and non-pregnancy shared similar microbial compositions when accounting for abundances (Fig 5d).

Finally, we examined the pregnant Lm-exposed cohort separately. While unsupervised clustering and analysis by Subject ID revealed no clusters (Fig 5e), weighted clustering indicated clustering by Subject (betadisper adj P = 0.002), but that the compositions were similar between subjects (Fig 5f). Furthermore, unsupervised clustering revealed distinct clustering of groups with and without APO (betadisper adj P = 0.018) with similar compositions between the two groups (Fig 5g). Weighted clustering indicated no significant clustering with occurrence of APO (Fig 5h).

To identify specific taxa that contributed to the differences indicated by our beta-diversity results, we performed non-parametric t-tests on the most abundant taxa associated with reproductive state and APO (Fig 6). We found four genera that significantly varied between pregnant and non-pregnant Lm-exposed subjects (Fig 6a). There was an increase in *Methanobrevibacter* spp. and *Treponema* spp., and a reduction in the *Eubacterium ruminantium* and *Prevotella* spp. in the pregnant subjects, compared to non-pregnant counterparts. There were two genera that significantly varied between those with or without APOs: an increase in *Akkermansia* spp. and a decrease in *Muribaculaceae* spp. (Fig 6b). It is worth noting that *Akkermansia* was only identifiable in two subjects (26 & 27).

We also examined the taxa in each subject of the pregnant Lm-exposed cohort (Fig 6c). While there were no significant associations with each subject within Cohort 4, we explored the OTUs with changes in abundance across the experimental period within each subject. We identified minor alterations within the *Treponema, Prevotella, Akkermansia, Lactobacillus, Lachnospiraceae, Streptococcus, Muribaculaceae, Eubacterium ruminantium* group*, Methanobrevibacter,* and *Phascolarctobacterium* (Fig 6c). *Akkermansia* also displayed significant changes within the same individual that was used in two pregnancy trials (subjects 26 & 27). Interestingly, *Akkermansia* was only identifiable within this individual. *Eubacterium ruminantium* had changes in abundance within subject 29, but not subsequent experimentation of that individual, subject 30. *Muribaculaceae* & *Methanobrevibacter* displayed minor changes among all subjects, with increased variability within subject 32 only. *Lachnospiraceae* displayed minor changes in abundance among all subjects, suggesting a potential interaction during gestational listeriosis. Furthermore, fewer changes to *Lachnospiraceae* in subject 27 as compared to subject 26 (same individual) indicate that *Lachnospira* may be more resistant to disruption with prior exposure. *Lactobacillus* and *Streptococcus* abundance changed primarily within subjects 25, 29, 32. *Phascolarctobacterium* had minor alterations in abundance for subjects 25, 27, 28, and 31. Interesting to note, variation within subject 27 was during that subject’s second experiment. *Prevotella* and *Treponema* have the largest range in abundances within subjects, indicating that these OTUs are highly susceptible to disruption during gestational listeriosis (Fig 6c). They were detected in all eight of the subjects in this cohort. In conclusion, alterations to low biomass genera including *Treponema, Prevotella, Akkermansia, Lactobacillus, Lachnospiraceae, Streptococcus, Muribaculaceae, Eubacterium ruminantium* group*, Methanobrevibacter,* and *Phascolarctobacterium* may be of particular interest in elucidating the associations of maternal GI microbes during gestational listeriosis.

To characterize the significant changes in abundance associated with reproductive state during listeriosis, as well as identify changes in low-abundance communities, we performed differential abundance analysis using the DESEQ package in R [36]. Differential analysis allows for comparing read counts between different conditions. We evaluated genera significantly associated with reproductive state, bacteremia, tissue bacterial burden, and APOs (Fig 7). In comparison of reproductive state, *Alloprevotella, Terrisporobacter, Rodentibacter, Actinobacillus, Romboutsia,* and *Agathobacter* were decreased, while *Bilophila* and RF39 were increased in the pregnant Lm-exposed cohort, compared to non-pregnant Lm-exposed subjects (Fig 7a). *Blautia, Helicobacter, Prevotellaceae_UCG-004, Akkermansia,* and *Lachnospiraceae_XPB1014*_group were all increased in subjects with bacteremia of the pregnant Lm-exposed cohort (Fig 7b). While *Lachnospira* was decreased, *Methanobrevibacter, Streptococcus, Marvinbryantia, Mogibacterium, p-1088-a5_gut_group, Intestinibacter, Enterorhabdus,* and *Akkermansia* were increased in pregnant Lm-exposed subjects with tissue infection with Lm (Fig 7c). Furthermore, in assessing genera with significant differences in abundance between those subject with APOs, only *Akkermansia* was identified as significantly increased in those with poor outcomes (Fig 7d).

**Figure 7(a-d).**
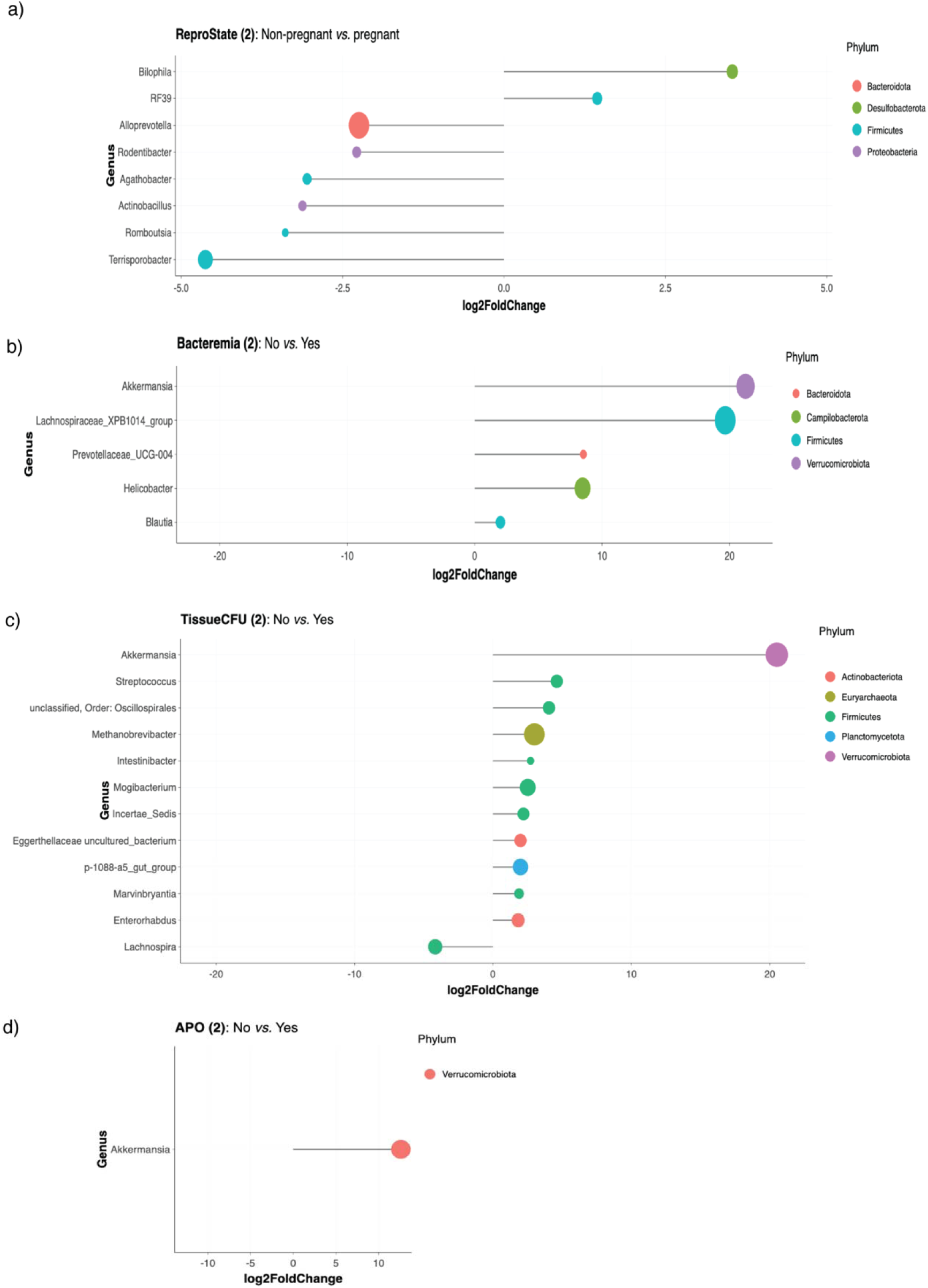
Differential Abundance Analysis of OTUs significantly associated with Reproductive State, Bacteremia, Tissue Positive for Lm, and APOs. The change in OTU abundance is presented on a log_2_ scale along the x-axis, and individual OTUs are listed on the left. Color indicates the phylum designation, and the size of the dot illustrates the relative abundance of that phylum within the whole population. Those data points to the right indicate an increase in abundance with the a) pregnant state, b) Lm bacteremia, c) presence of tissue Lm, and d) occurrence of an APO, while negative values indicating a decrease in the specific OTU associated with these parameters. Please note the scale of the x-axis varies by graph. The main findings from this analysis are summarized below (Table 1).

**Table 1.**
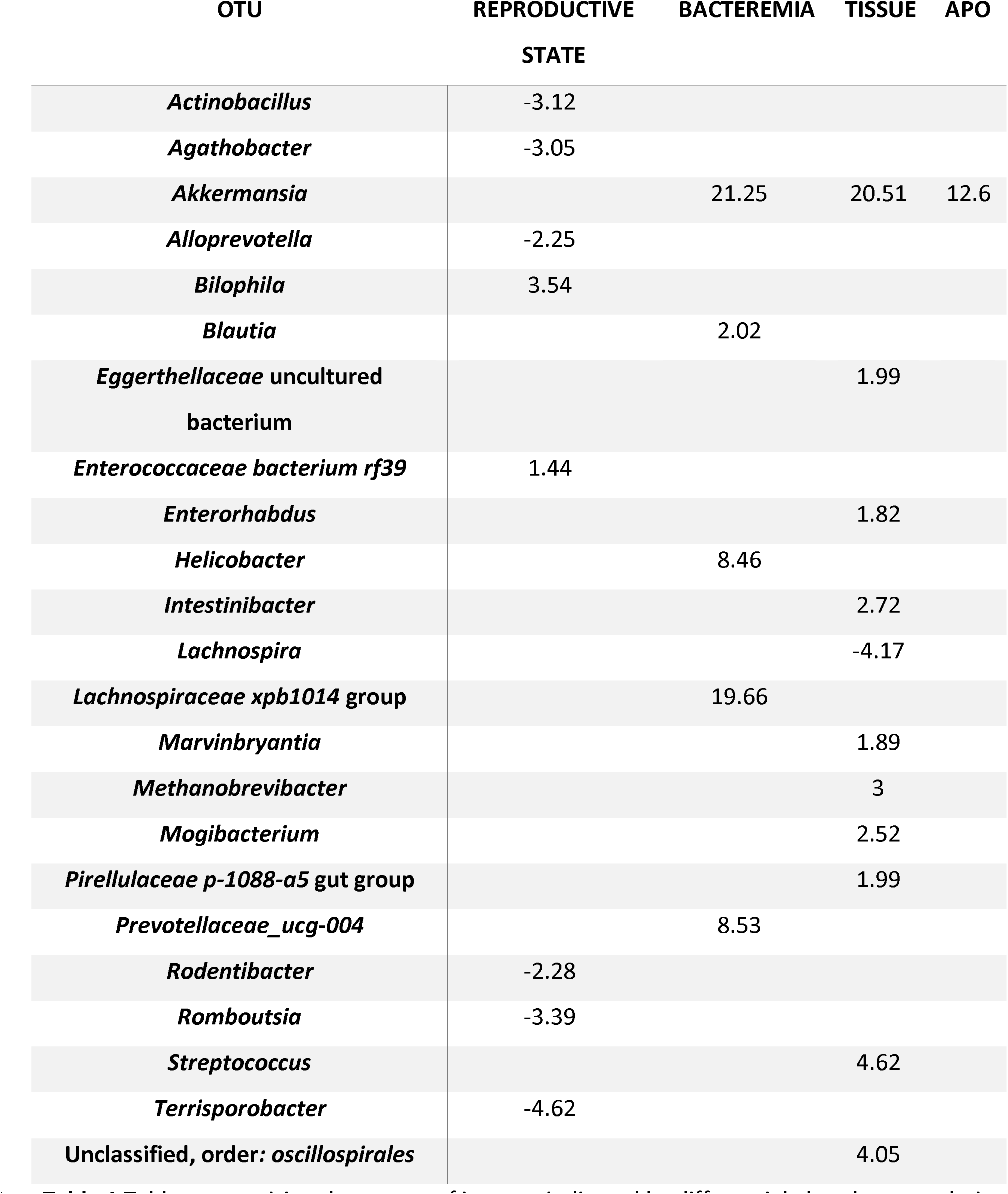
Table summarizing the genera of interest indicated by differential abundance analysis first by reproductive state in Lm-exposed Cohorts 2 and 4 (n= 58), then by incidence of bacteremia, bacterial burden, and adverse pregnancy outcome within Lm-exposed pregnant Cohort 4 only (n= 29). The increased or decreased presence of each OTU associated with each factor is indicated as log_2_-fold change.

From our comparisons, we found 23 OTUs whose abundance was significantly associated with reproductive state, Lm bacteremia, Lm infection of the tissues, and APO during gestational listeriosis (Supplemental Table 4). For ease of the reader, the main findings are summarized below (Table 1).

## Discussion

Evidence in the literature supporting microbial interactions during pathogen exposure led us to evaluate the potential impact of listeriosis on maternal GI microbial communities in pregnant and nonpregnant subjects. Contrary to our hypothesis, there was no significant change in GI community richness or abundance associated solely with exposure to Lm in nonpregnant animals. However, in the case of gestational listeriosis, we identified significant remodeling to genera including *Eubacterium ruminantium, Methanobrevibacter, Prevotella,* and *Treponema*. Our findings further indicate an association of the maternal intestinal commensal microbes with the pathogenesis of listeriosis during pregnancy.

Lm is a Gram-positive organism, with thick outer layer consisting of a layer of dense peptidoglycan, enabling the bacterium to survive and replicate across a wide range of temperatures, pH and salt concentrations[37]. These traits enable Lm to withstand the highly acidic environment of the stomach, as well as bile within the gallbladder where replication occurs [37, 38]. Further in the gastrointestinal tract, Lm invades host cells, enters the enterocytes and goblet cells in the small intestine, cecum, and colon, and gains access to the lymphatic system through a process known as paracytosis, and ultimately enters the bloodstream[39, 40]. Lm virulence genes that facilitate host cell invasion include the bacterial surface proteins internalin A (InlA) and internalin B (InlB) [40]. Within the host cell, Lm secretes Listeriolysin O (LLO) along with phospholipases PlcA and PlcB to escape from vacuoles into the cytosol, where the bacterium can replicate [39]. Bacterial surface proteins ActA and PrfA promote cell-to-cell spread, furthering infection and evading extracellular immune detection [37]. More recently characterized, InlP has been shown to interact with affadin to invade cells at the MFI [10–12]. Each of these virulence factors aid Lm in invasion, survival, and replication within the host cells.

Once within the circulatory system, Lm disseminates to the liver, spleen, gallbladder, and the placenta[41, 42]. Within the intervillous space of the placenta, exchange of nutrients from mother to fetus occurs, including amino acids, fatty acids, glucose, and oxygen to underpin fetal development [8]. Through incompletely defined mechanisms, Lm is able to attach to and invade the placental tissues [11, 12]. Once within the placental tissue, Lm may establish severe infection which ultimately causes acute inflammation, chorioamnionitis, and necrosis[9]. Within the gastrointestinal (GI) tract, there are two barriers to infection: mechanical, which consists of epithelial enterocytes and an associated layer of mucus, and environmental, which consists of immune cells, cytokines, metabolites, hormones, and microorganisms [43]. The potential impact of pregnancy on these barriers is poorly understood.

Interactions between GI bacteria and the host can benefit the host through the modulation of nutrient uptake and metabolism, strengthening the intestinal barrier function, inhibiting pathogen propagation, and regulating host immunity [44–46]. This communication occurs via bacterial metabolic products such as the short-chain fatty acids (SCFA) propionate, butyrate, acetate, formate, and succinate that are produced by degradation and fermentation of dietary fiber, vitamins, and immunomodulatory peptides [47, 48]. SCFAs are the most extensively studied bacterial metabolic pathways in the context of host immunity [49] and are suggested to play a pivotal role in host-microbial crosstalk [50]. SCFAs have been shown to improve epithelial barrier function and immune tolerance and to promotes gut homeostasis [51, 52]. However, the impact of SCFA specifically on Lm remains unknown. Furthermore, SCFAs have been show to increase mucus production by stimulating epithelial mucin-2 expression [50]. Butyrate, in particular, plays a critical role in energy intestinal motility, immunomodulation, suppression of inflammation in the gut, and has been further shown to inhibit production of virulence factors in Lm [53]. There is ample evidence that commensal microorganisms confer protection against invading pathogens, and defense against Lm invasion, potentially through the production of SCFAs [19, 54–56]. It is thus important to consider whether the current results provide any insight into whether alterations in the gut microbiota may be complicit in the susceptibility of pregnancy to disseminated listeriosis.

Our study identified four genera that vary significantly across all timepoints between pregnant and non-pregnant Lm-exposed subjects (Fig 6a). There was an increase in relative abundance within the entire population of *Methanobrevibacter* spp. and *Treponema* spp., but a reduction in the *Eubacterium ruminantium* group and *Prevotella* spp. within in the pregnant subjects, compared to non-pregnant counterparts. However, when examining the differential abundance between reproductive states, we identified decreases in *Alloprevotella, Terrisporobacter, Rodentibacter, Actinobacillus, Romboutsia,* and *Agathobacter*, and an increase in *Bilophila* and RF39 (Fig 7a).

*Methanobrevibacter* is strictly anaerobic archaebacteria that produces methane through the reduction of carbon dioxide via hydrogen. One study in human pregnancy showed that this genus was differentially abundant between those with zero or high parity, or the number of times a person has given birth [15]. Moreover, that study also showed that as parity increases, microbial remodeling occurs more rapidly [15]. The significance of changes in these taxa remains to be elaborated. Our data further supports that *Methanobrevibacter* is impacted by reproductive state, as seen by an increase in abundance associated with the pregnant cohorts.

*Treponema*, a member of the phyla Spirochaetota, contains species known to cause syphilis and yaws in humans and genital ulcers in baboons [57]. It has been documented that *Treponema* is a naturally occurring infection in primates, with extensive studies using NHP experimental models of Treponematoses [58]. During pregnancy, infection with *T. pallidum* can lead to early fetal loss, preterm birth, stillbirth, low birth weight, and congenital disease [59]. In the context of listeriosis, there is some evidence that intravenous *T. pallidum* is associated with resistance to intravenous listeria infection [60]. Although the exact mechanisms of this potential interaction are undefined, it is hypothesized that *T. pallidum* triggers cell mediated immunity which prolongs the listericidal activity[61].

*Eubacterium ruminantium* is a Gram-positive bacterium that plays pivotal role in metabolism, producing methane, butyrate, lactate, and formate [62]. The genus *Eubacteria*, belonging to the phylum Firmicutes, includes a myriad of diverse species that have potential as therapeutic microbes. Although it is a commonly documented in the human gut microbiome, most of the knowledge about this genus originates from ruminant microbiome studies [63–65].

*Prevotellaceae* is a predominant taxon in the rhesus monkey gut [66] and has been documented as one of the most abundant taxa within the human gut [21]. One study that examined age-associated microbial communities in mice found that Lm infection increased the abundance of *Prevotellaceae* in young-adult mice [67]. Another study found enrichment of *Prevotella* relative to *Listeria* [68]. Our data support these findings, as we identified an increase in the *Prevotellaceae* associated with bacteremia (Fig 5b).

While there were no significant changes in the relative abundances of top 10 abundant taxa associated with bacteremia or tissue infection with Lm, we were able to identify alterations in less predominant genera. *Blautia, Helicobacter, Prevotellaceae_UCG-004, Akkermansia* and *Lachnospiraceae_XPB1014_group* were all increased in animals with bacteremia (Fig 7b). While *Methanobrevibacter, Streptococcus, Marvinbryantia, Mogibacterium, p-1088-a5_gut_group, Intestinibacter, Enterorhabdus, and Akkermansia* were increased in animals with tissue infection of Lm, *Lachnospira* was decreased (Fig 7c). While *Lachnospiraceae* was increased with bacteremia, it was decreased with tissue infection of Lm. The impact of *Lachnospiraceae* on the host physiology is inconsistent across different studies [69]. This genus has been associated with various intra- and extra- intestinal diseases [69]. Members of the *Lachnospiraceae* family were shown to be significantly increased in aged mice with listeriosis [67]. It is important to note that members of the *Lachnospiraceae* spp. include some of the most prolific producers of SCFA, a microbial byproduct as discussed above.

There were marked changes to both *Akkermansia* spp. and *Muribaculaceae* spp. in macaques with APOs (Fig. 7b). Through examining differential abundance, we identified a significant increase of *Akkermansia* in those with poor outcomes (Fig 7d). It is interesting to note that *Akkermansia* was only identifiable at low levels within 2 subjects, the same individual who was utilized twice in the experiment (26 & 27). There is growing interest in *Akkermansia* due to its potential association with intestinal health. Notably, reduced levels of *A. muciniphila* have been observed in patients with inflammatory bowel diseases and metabolic disorders, suggesting it may have potential anti-inflammatory properties [70]. One listeriosis study examined age-associated microbial communities in mice and found that *Akkermansia* were only abundant in infected young-adult mice, with diminished abundance in infected aged mice[67]. While the subjects utilized in this study were of reproductive age, it is possible that individual 26/27 may have had some circumstances facilitating *Akkermansia* colonization/prevalence. Alternatively, it may have been present in other subjects, but at levels not detectable by 16S rRNA sequencing.

The changes in gut microbiota may point to potential alternatives to antibiotic treatment in pregnancy. Epidemiological studies also have found an association between antibiotic usage during pregnancy and increased incidence of asthma in the infant [71, 72]. While antibiotics can treat listeriosis, the risks associated with this treatment leave clinicians and patients desiring safer alternatives such as preventative biotherapies [73, 74]. One potential treatment is probiotic intervention, either as a daily preventative or through microbial transplant during severe listeriosis [75–77]. Probiotic intervention is clinically used to treat patients with chronic bowel disease, ulcerative colitis, and necrotizing enterocolitis [76]. Bacterial genera commonly utilized in probiotic treatments include, but are not limited to *Lactobacterium, Bacillis, Bifidobacterium, Bacteriodes, and Akkermansia* [78]. A review of probiotic administration as a potential maternity supplement highlighted the importance of understanding microbial interactions during pregnancy and their potential impact on reproductive health outcome [79].

In summary, we identified genera whose abundances are linked with reproductive state, bacteremia, tissue infection, and APO during listeriosis. Of the 23 OTUs of interest that were significantly associated with gestational listeriosis and disease progression (Fig 7), our data indicate that *Treponema*, *Prevotella*, *Akkermansia*, *Eubacterium ruminantium* group, and *Methanobrevibacter* are key genera in understanding the influences of and on the maternal gastrointestinal microbiota in susceptibility to listeriosis.

## Conclusions

These findings indicate that dysbiosis is not associated with reproductive state or listeriosis alone. Dysbiosis is significantly associated with the interaction of listeriosis during pregnancy, bolstering the clinical significance of increased infection susceptibility in human gestational listeriosis. This implies that exposure to listeriosis exacerbates the mild disruption that may be associated with the pregnant state. Our data reinforce the previous notion that the pregnant state is uniquely susceptible to listeriosis and builds on our understanding of the potential role of the microbiome in maternal-fetal health through identification of OTUs of primary interest. Further investigation to characterize the gut microbial environment during gestation may provide insight into treating listeriosis during pregnancy.

## Acknowledgements

We thank the WNPRC Veterinary, Scientific Protocol Implementation, Colony Services, and Pathology Services staff for assistance with animal procedures, including breeding, ultrasound monitoring, and sample collection. We thank Sophia Kathariou of North Carolina State University for generous donation of clinical strain LM2203. We would like to thank Faye Hartmann and the Clinical Pathology Laboratory at the School of Veterinary Medicine for assistance with blood cultures, and Greg Wiepz at the WNPRC for assistance with specimen processing. We sincerely thank Bryce Wolfe for assistance with study design, sample collection, and mentorship.

