## Supplemental Figures & Tables for "*Listeria monocytogenes* infection in pregnant macaques alters the maternal gut microbiome"

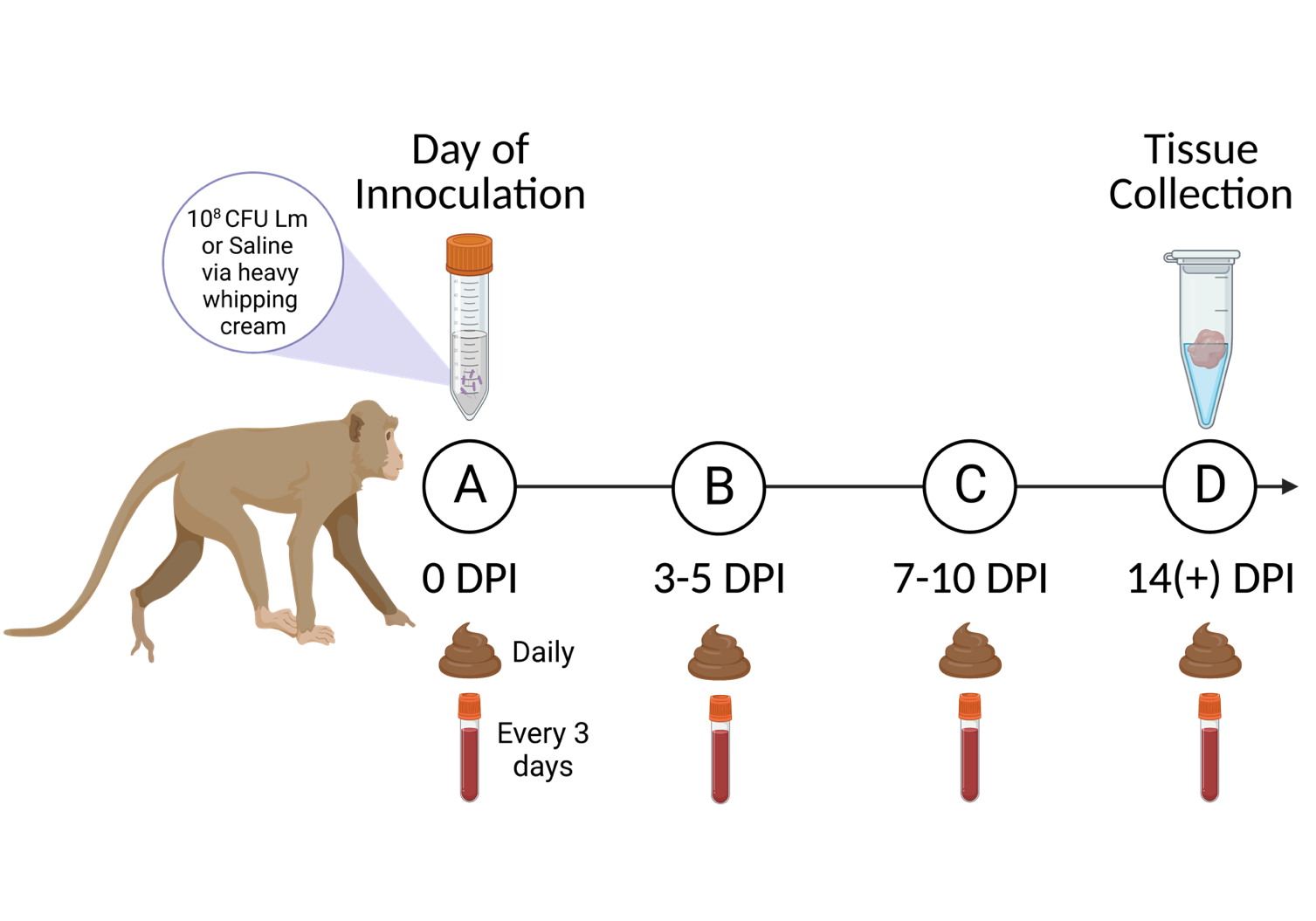

**Supplemental Figure 1 Collection Timeline**

The image above depicts the collection of fecal and blood samples at timepoints 0dpi (A), 3-5 dpi (B), 7-10 dpi (C), and 14+ dpi (D). The animal model received 10^8^ CFU of Lm or saline control via heavy whipping cream at timepoint A. Fecal and blood samples were collected prior to administration of whipping cream, and at regular intervals (daily collection of feces, blood samples every 3 days) until tissue collection.

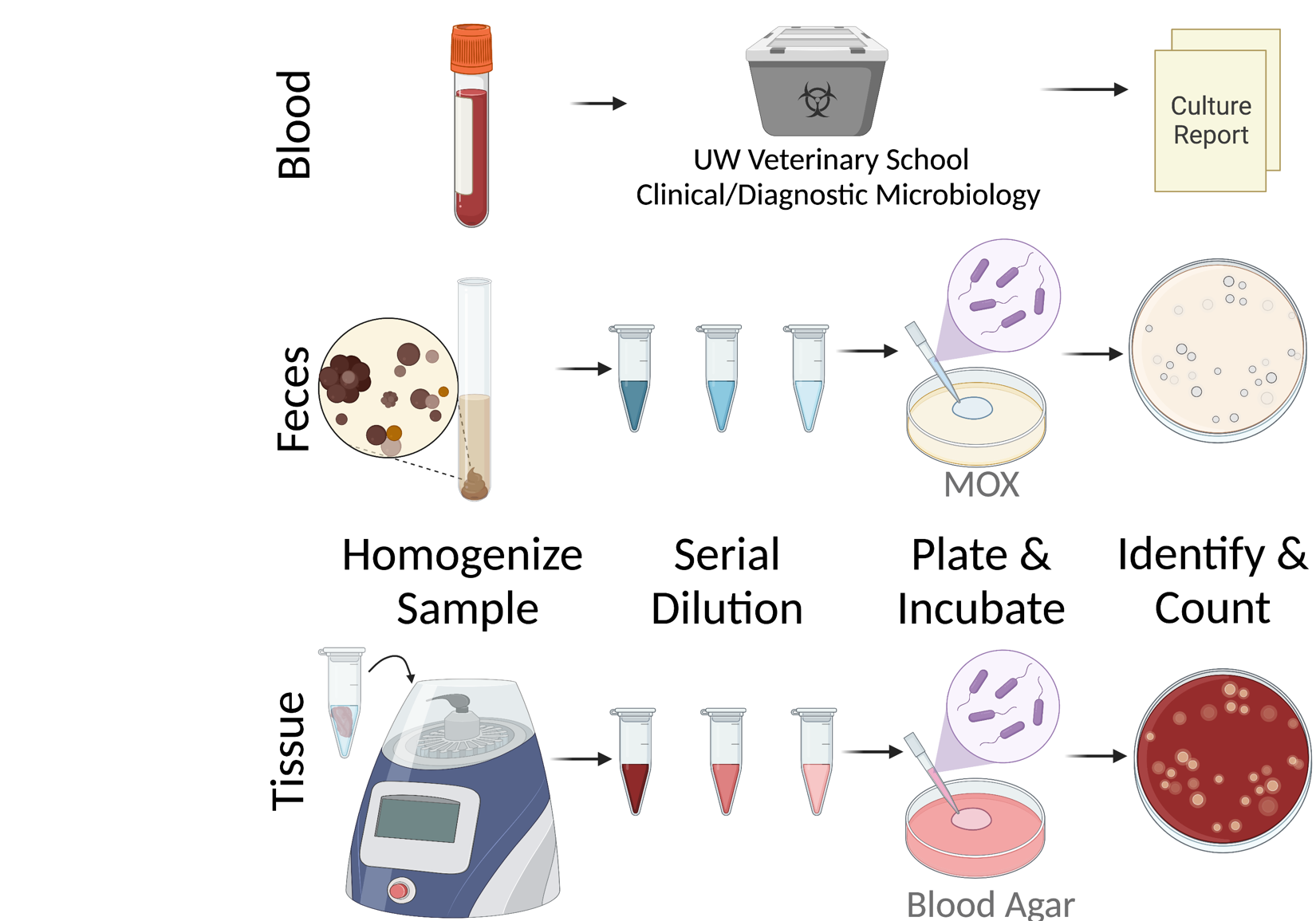

**Supplementary Figure 2** Summary of the sample processing and method of identification of Lm. Blood samples were submitted to UW Veterinary School Clinical Diagnostic Microbiology for mass spectrometry identification of bacterial pathogens. Fecal samples were serially diluted and plated on Modified Oxford Medium for incubation and counting. Tissue samples were homogenized, serially diluted, and plated on blood agar for incubation and counting.

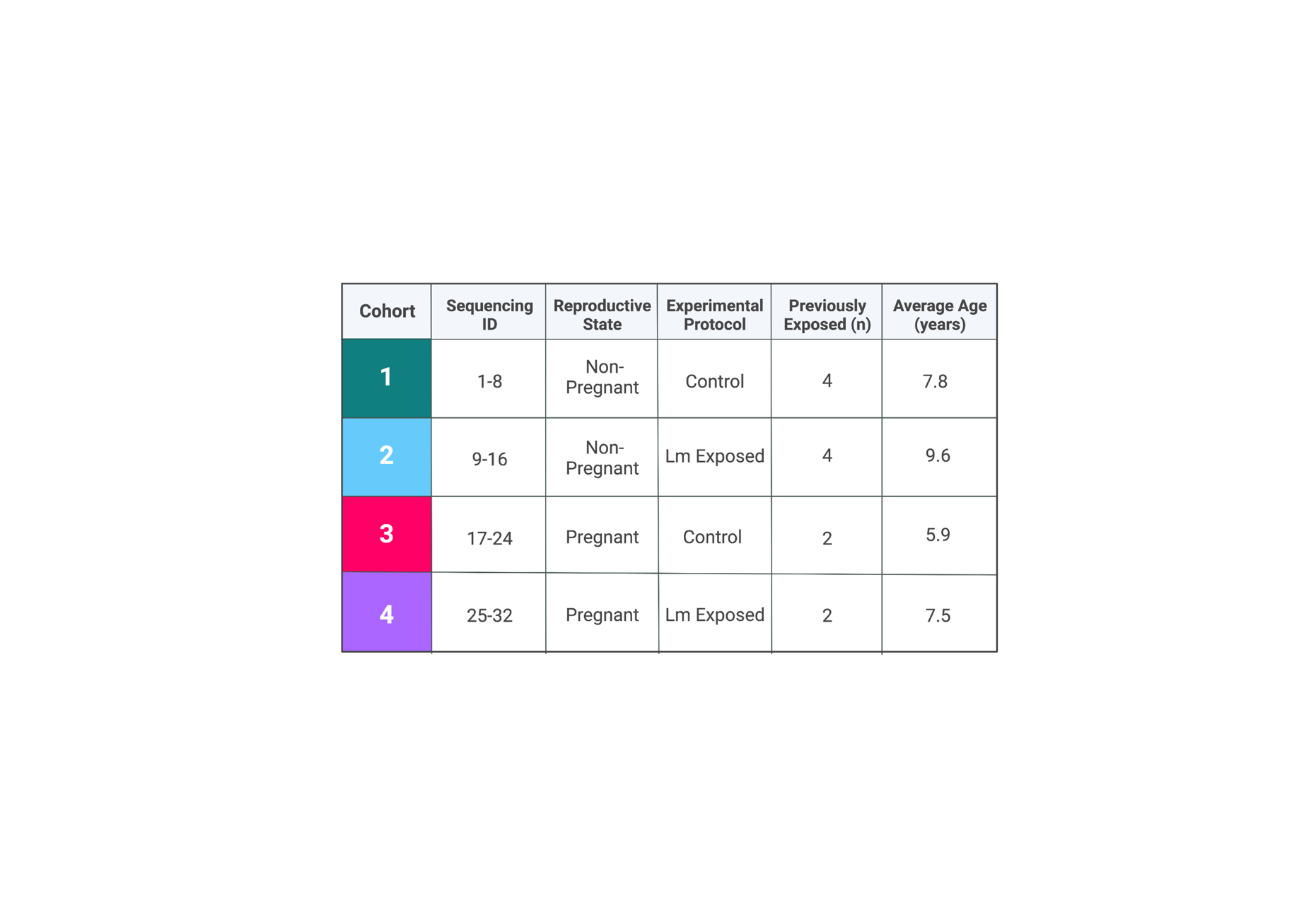

**Supplementary Table 1** Experimental cohorts are presented, coded by color, along with their respective sequencing IDs, Reproductive State, Experimental Protocol, number of those previously exposed to Lm, and the average age of the subjects for each cohort.

| **1 & 2** | **3 & 4** | **All Cohorts** |
| --- | --- | --- |
| Colon | Placenta | Liver |
| Cecum | Decidua | Spleen |
| Jejunum | Fetal Tissues | Lymph Node |
| Endometrium |  |  |

**Supplementary Table 2** Summary of the tissues collected for each cohort. All cohorts had liver, spleen, and lymph node collected for subsequent analysis. However, due to study limitations, colon, cecum, jejunum, and endometrium were collected from terminal subjects only, the non-pregnant cohorts. Furthermore, placenta, decidua, and fetal tissues were only collected from the pregnant cohorts.

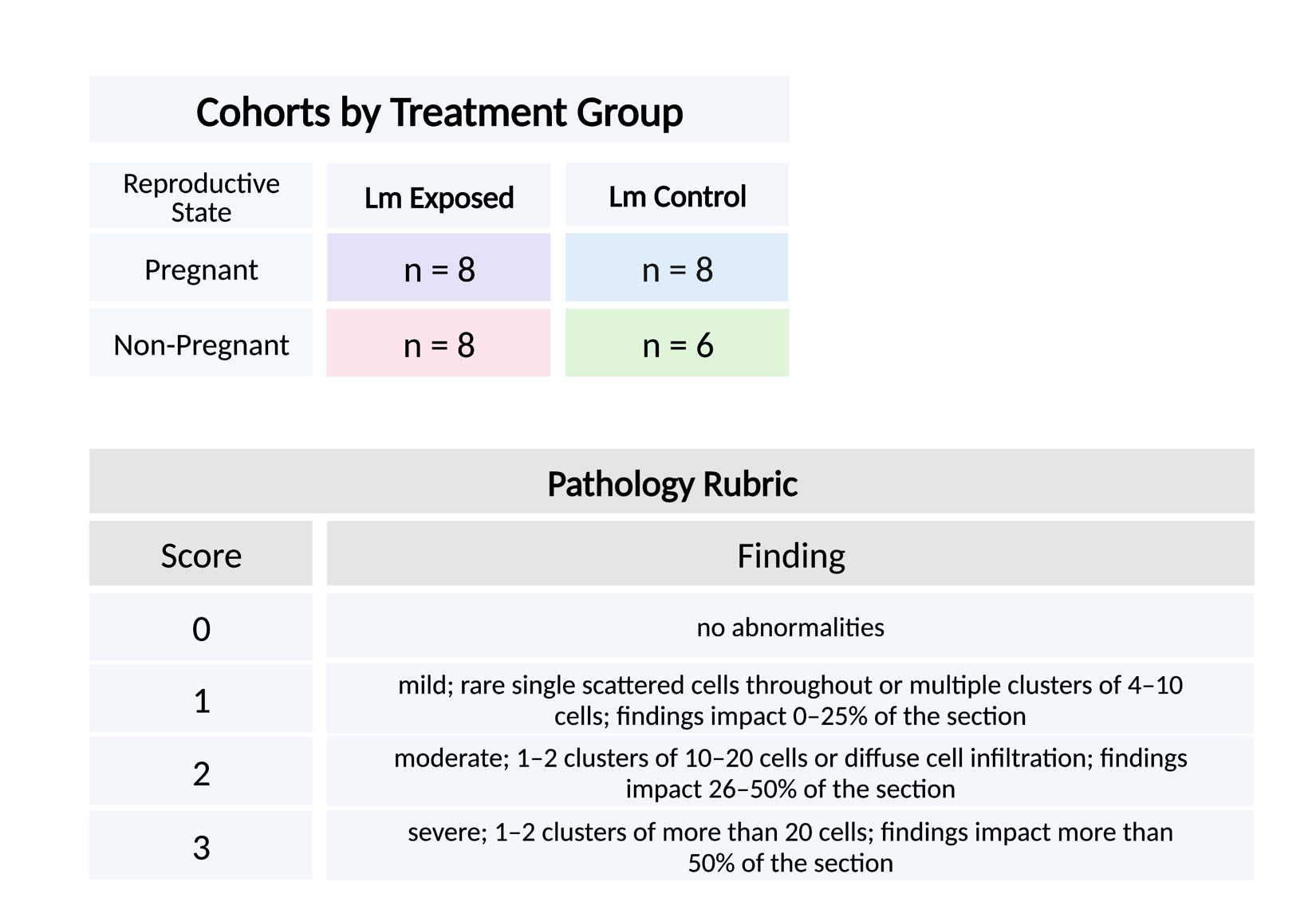
**Supplementary Table 3** Scoring rubric used for the tissue histopathology.

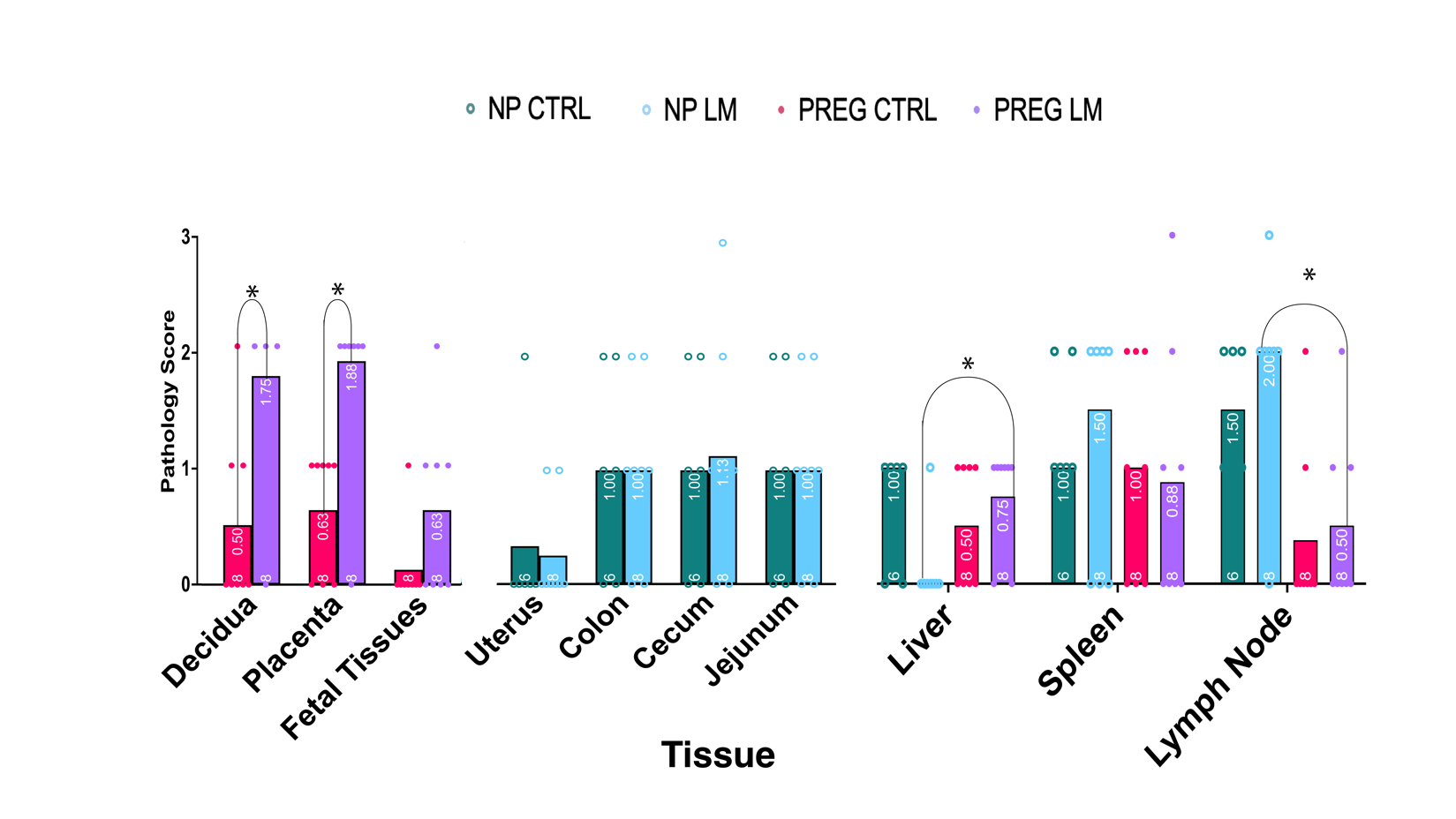

**Supplementary Figure 3** The Average Tissue Pathology Score for each tissue collected, by cohort. Each individual score is plotted with a circle, the n listed at the bottom of the bar, and the average value listed at the top of the bar. Nonpregnant cohorts are depicted by open circles, pregnant by closed circles. Significance is denoted with an asterisk and the significantly different tissues are connected by lines. * = P ≤ 0.05 and ** = P ≤ 0.01

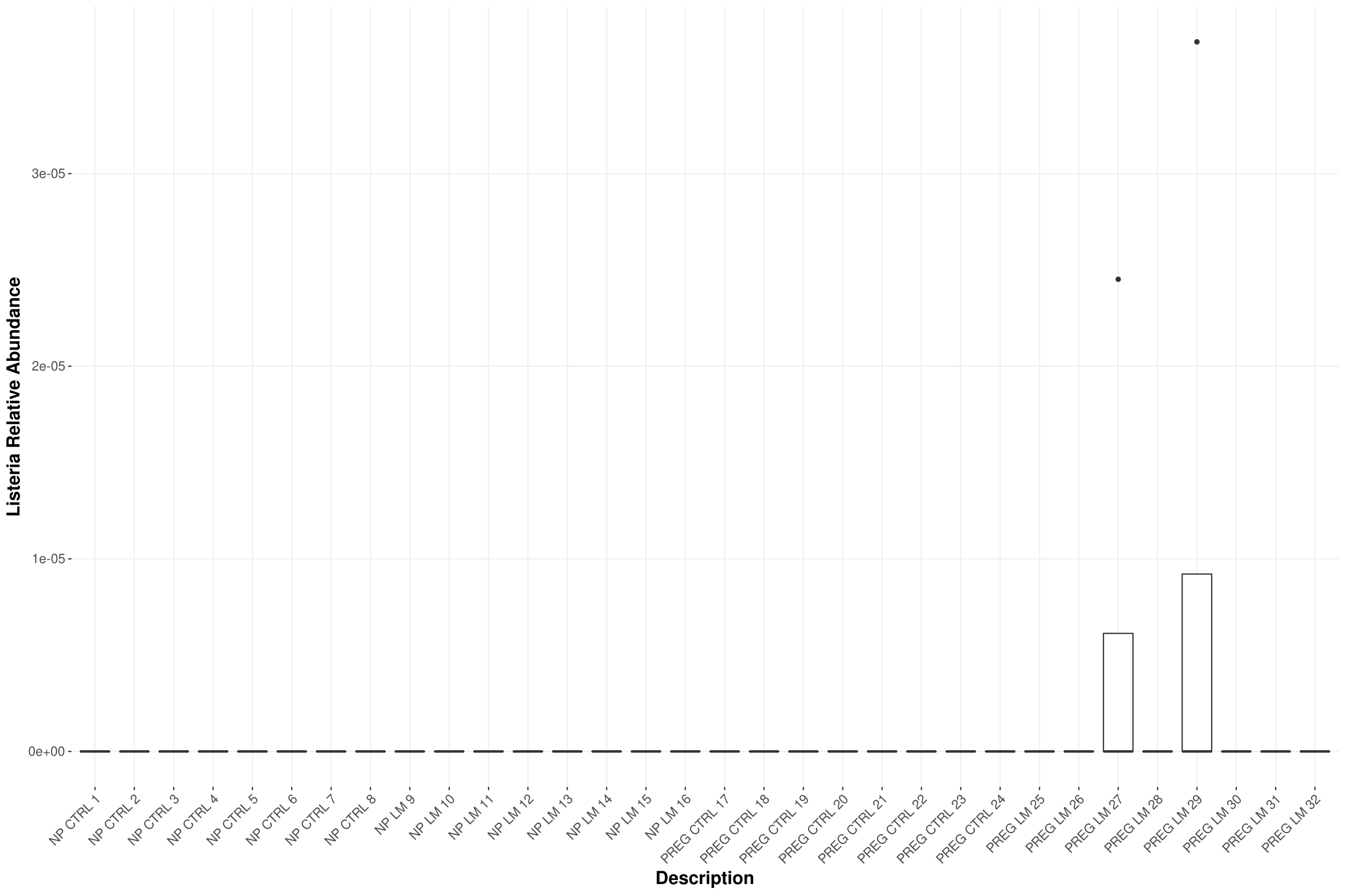
 **Supplementary Figure 4** Abundance of the *Listeria monocytogenes* taxon identified per individual. Only 2 individuals from the Pregnant Lm-exposed cohorts had Lm identified by 16S sequencing and analysis.

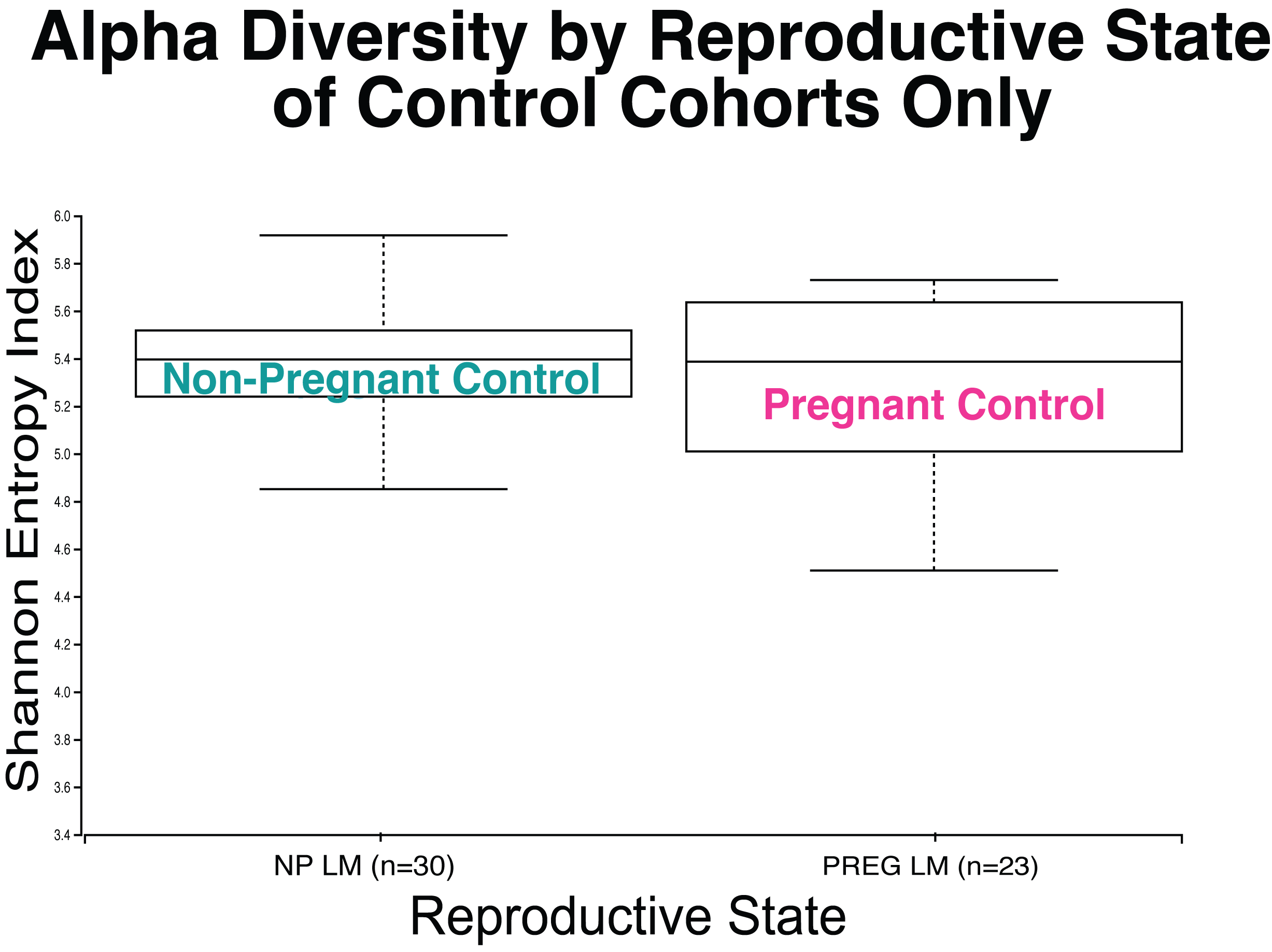

**Supplementary Figure 5** Alpha Diversity by Reproductive State of Control Cohorts. The box plot shows the H-value of the Shannon Entropy Index, a measure of α-diversity on the y-axis, with bars indicating SEM. The x-axis shows Cohort 1 (non-pregnant controls, n=) compared to Cohort 3 (pregnant controls, n=23).

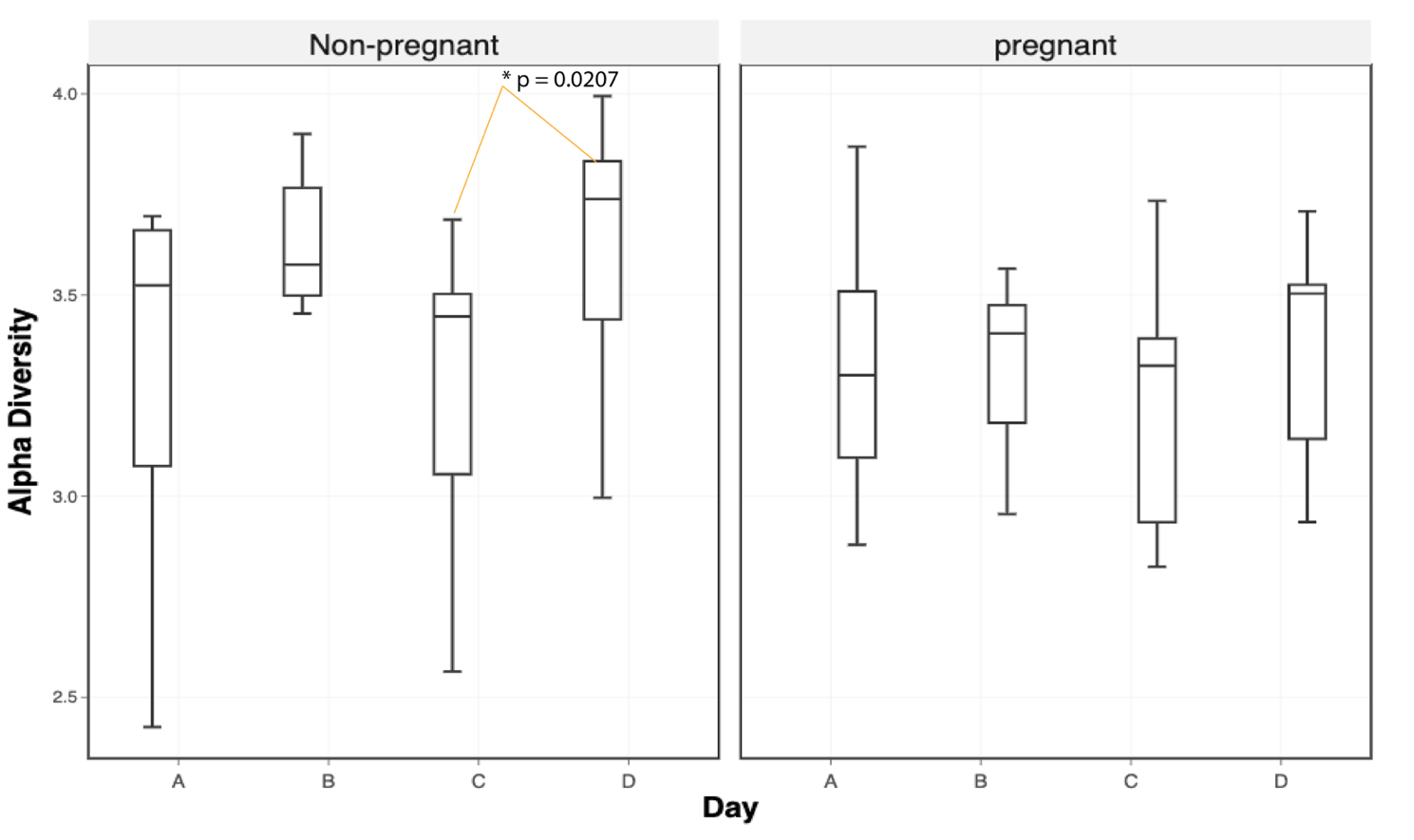

**Supplementary Figure 6.** Alpha Diversity by Day. The box plots indicate the alpha diversity values of Cohort 2 (non-pregnant Lm-exposed, n= 30) and Cohort 4 (pregnant Lm-exposed, n=23) during each collection timepoints ( A: 0dpi, B: 3-5 dpi, C: 7-20 dpi, and D: 14+ dpi). The bars indicate SEM. Significance denoted by asterisks and the significantly different factors are connected by yellow lines. * P ≤ 0.05.

**Supplementary Table 4 – Statistical Analyses**

| **Figure 1** |  |  |  |  |  |  |
| --- | --- | --- | --- | --- | --- | --- |
| **1a) Weight** |  |  |  |  |  |  |
| Table Analyzed | Copy of Weight | |  |  |  |  |
| Data sets analyzed | A-D |  |  |  |  |  |
| One- Way ANOVA summary |  |  |  |  |  |  |
| F | 0.1545 |  |  |  |  |  |
| P value | 0.9259 |  |  |  |  |  |
| P value summary | ns |  |  |  |  |  |
| Significant diff. among means (P < 0.05)? | No |  |  |  |  |  |
| R squared | 0.01628 |  |  |  |  |  |
| Brown-Forsythe test |  |  |  |  |  |  |
| F (DFn, DFd) | 1.019 (3, 28) |  |  |  |  |  |
| P value | 0.3991 |  |  |  |  |  |
| P value summary | ns |  |  |  |  |  |
| Are SDs significantly different (P < 0.05)? | No |  |  |  |  |  |
| Bartlett's test |  |  |  |  |  |  |
| Bartlett's statistic (corrected) | 6.615 |  |  |  |  |  |
| P value | 0.0852 |  |  |  |  |  |
| P value summary | ns |  |  |  |  |  |
| Are SDs significantly different (P < 0.05)? | No |  |  |  |  |  |
| ANOVA table | SS | DF | MS | F (DFn,DFd) | P value |  |
| Treatment (between columns) | 0.05663 | 3 | 0.01888 | F (3, 28) = 0.1545 | P=0.9259 |  |
| Residual (within columns) | 3.421 | 28 | 0.1222 |  |  |  |
| Total | 3.478 | 31 |  |  |  |  |
| Data summary |  |  |  |  |  |  |
| Number of treatments (columns) | 4 |  |  |  |  |  |
| Number of values (total) | 32 |  |  |  |  |  |
| Multiple Comparisons |  |  |  |  |  |  |
| Number of families | 1 |  |  |  |  |  |
| Number of comparisons per family | 6 |  |  |  |  |  |
| Alpha | 0.05 |  |  |  |  |  |
| Tukey's multiple comparisons test | Mean Diff. | 95.00% CI of diff. | Below threshold? | Summary | Adjusted P Value | |
| NP Ctrl vs. NP Lm | -0.027 | -0.5042 to 0.4502 | No | ns | 0.9987 | A-B |
| NP Ctrl vs. Preg Ctrl | -0.007 | -0.4842 to 0.4702 | No | ns | >0.9999 | A-C |
| NP Ctrl vs. Preg Lm | -0.1058 | -0.5830 to 0.3715 | No | ns | 0.9296 | A-D |
| NP Lm vs. Preg Ctrl | 0.02 | -0.4572 to 0.4972 | No | ns | 0.9995 | B-C |
| NP Lm vs. Preg Lm | -0.07875 | -0.5560 to 0.3985 | No | ns | 0.969 | B-D |
| Preg Ctrl vs. Preg Lm | -0.09875 | -0.5760 to 0.3785 | No | ns | 0.9416 | C-D |
| Test details | Mean 1 | Mean 2 | Mean Diff. | SE of diff. | n1 | n2 |
| NP Ctrl vs. NP Lm | -0.237 | -0.21 | -0.027 | 0.1748 | 8 | 8 |
| NP Ctrl vs. Preg Ctrl | -0.237 | -0.23 | -0.007 | 0.1748 | 8 | 8 |
| NP Ctrl vs. Preg Lm | -0.237 | -0.1313 | -0.1058 | 0.1748 | 8 | 8 |
| NP Lm vs. Preg Ctrl | -0.21 | -0.23 | 0.02 | 0.1748 | 8 | 8 |
| NP Lm vs. Preg Lm | -0.21 | -0.1313 | -0.07875 | 0.1748 | 8 | 8 |
| Preg Ctrl vs. Preg Lm | -0.23 | -0.1313 | -0.09875 | 0.1748 | 8 | 8 |
| **1b) Diarrhea of All Cohorts** |  |  |  |  |  |  |
| Table Analyzed | Diarhhea Data | |  |  |  |  |
| Three-way ANOVA | Matching by factors: REPROSTATE & LMEXPOSURE | | | |  |  |
| Assume sphericity? | Yes |  |  |  |  |  |
| Alpha | 0.05 |  |  |  |  |  |
| Source of Variation | % of total variation | P value | P value summary | Significant? |  |  |
| TIME | 3.925 | 0.2744 | ns | No |  |  |
| REPROSTATE | 2.395 | 0.1272 | ns | No |  |  |
| LMEXPOSURE | 2.395 | 0.0608 | ns | No |  |  |
| TIME x REPROSTATE | 1.331 | 0.7141 | ns | No |  |  |
| TIME x LMEXPOSURE | 2.395 | 0.3029 | ns | No |  |  |
| REPROSTATE x LMEXPOSURE | 1.663 | 0.0533 | ns | No |  |  |
| TIME x REPROSTATE x LMEXPOSURE | 2.861 | 0.0955 | ns | No |  |  |
| SUBJECTSCORE | 26.88 |  |  |  |  |  |
| SUBJECTSCORE x REPROSTATE | 27.15 |  |  |  |  |  |
| SUBJECTSCORE x LMEXPOSURE | 17.56 |  |  |  |  |  |
| ANOVA table | SS | DF | MS | F (DFn, DFd) | P value |  |
| TIME | 1.844 | 3 | 0.6146 | F (3, 28) = 1.363 | P=0.2744 |  |
| REPROSTATE | 1.125 | 1 | 1.125 | F (1, 28) = 2.471 | P=0.1272 |  |
| LMEXPOSURE | 1.125 | 1 | 1.125 | F (1, 28) = 3.818 | P=0.0608 |  |
| TIME x REPROSTATE | 0.625 | 3 | 0.2083 | F (3, 28) = 0.4575 | P=0.7141 |  |
| TIME x LMEXPOSURE | 1.125 | 3 | 0.375 | F (3, 28) = 1.273 | P=0.3029 |  |
| REPROSTATE x LMEXPOSURE | 0.7813 | 1 | 0.7813 | F (1, 28) = 4.070 | P=0.0533 |  |
| TIME x REPROSTATE x LMEXPOSURE | 1.344 | 3 | 0.4479 | F (3, 28) = 2.333 | P=0.0955 |  |
| SUBJECTSCORE | 12.63 | 28 | 0.4509 |  |  |  |
| SUBJECTSCORE x REPROSTATE | 12.75 | 28 | 0.4554 |  |  |  |
| SUBJECTSCORE x LMEXPOSURE | 8.25 | 28 | 0.2946 |  |  |  |
| Residual | 5.375 | 28 | 0.192 |  |  |  |
| Data summary |  |  |  |  |  |  |
| Number of columns | 2 x 2 |  |  |  |  |  |
| Number of rows (TIME) | 4 |  |  |  |  |  |
| Number of subjects (SUBJECTSCORE) | 32 |  |  |  |  |  |
| Number of missing values | 0 |  |  |  |  |  |
| Multiple Comparisons |  |  |  |  |  |  |
| Compare row means (main row effect) |  |  |  |  |  |  |
| Number of families | 1 |  |  |  |  |  |
| Number of comparisons per family | 6 |  |  |  |  |  |
| Alpha | 0.05 |  |  |  |  |  |
| Tukey's multiple comparisons test | Mean Diff. | 95.00% CI of diff. | Below threshold? | Summary | Adjusted P Value | |
| A vs. B | -0.3125 | -0.7708 to 0.1458 | No | ns | 0.2672 |  |
| A vs. C | -0.25 | -0.7083 to 0.2083 | No | ns | 0.4571 |  |
| A vs. D | -0.25 | -0.7083 to 0.2083 | No | ns | 0.4571 |  |
| B vs. C | 0.0625 | -0.3958 to 0.5208 | No | ns | 0.982 |  |
| B vs. D | 0.0625 | -0.3958 to 0.5208 | No | ns | 0.982 |  |
| C vs. D | 0 | -0.4583 to 0.4583 | No | ns | >0.9999 |  |
| Test details | Mean 1 | Mean 2 | Mean Diff. | SE of diff. | N1 | N2 |
| A vs. B | 0.0625 | 0.375 | -0.3125 | 0.1679 | 32 | 32 |
| A vs. C | 0.0625 | 0.3125 | -0.25 | 0.1679 | 32 | 32 |
| A vs. D | 0.0625 | 0.3125 | -0.25 | 0.1679 | 32 | 32 |
| B vs. C | 0.375 | 0.3125 | 0.0625 | 0.1679 | 32 | 32 |
| B vs. D | 0.375 | 0.3125 | 0.0625 | 0.1679 | 32 | 32 |
| C vs. D | 0.3125 | 0.3125 | 0 | 0.1679 | 32 | 32 |
| **1d) Diarrhea by Subject of Cohort 4** |  |  |  |  |  |  |
| Table Analyzed | Preg Lm Diarhhea | |  |  |  |  |
| Two-way ANOVA | Ordinary |  |  |  |  |  |
| Alpha | 0.05 |  |  |  |  |  |
| Source of Variation | % of total variation | P value | P value summary | Significant? |  |  |
| Day | 10.67 | 0.2599 | ns | No |  |  |
| Subject | 37.41 | 0.0812 | ns | No |  |  |
| ANOVA table | SS (Type III) | DF | MS | F (DFn, DFd) | P value |  |
| Day | 2.344 | 3 | 0.7813 | F (3, 21) = 1.438 | P=0.2599 |  |
| Subject | 8.219 | 7 | 1.174 | F (7, 21) = 2.162 | P=0.0812 |  |
| Residual | 11.41 | 21 | 0.5432 |  |  |  |
| Data summary |  |  |  |  |  |  |
| Number of columns (Subject) | 8 |  |  |  |  |  |
| Number of rows (Day) | 4 |  |  |  |  |  |
| Number of values | 32 |  |  |  |  |  |
| **Figure 5 - Beta-Diversity Statistics performed in R** | |  |  |  |  |  |
| **5a) Bray Treatment group** |  |  |  |  |  |  |
| adonis2(DISTWAR ~ TreatmentGroup, data = FUCREA) | |  |  |  |  |  |
| Permutation test for adonis under reduced model | |  |  |  |  |  |
| Terms added sequentially (first to last) |  |  |  |  |  |  |
| Permutation: free |  |  |  |  |  |  |
| Number of permutations: 999 |  |  |  |  |  |  |
| adonis2(formula = DISTWAR ~ TreatmentGroup, data = FUCREA) | | |  |  |  |  |
| Df SumOfSqs      R2      F Pr(>F) | |  |  |  |  |  |
| TreatmentGroup   3   1.7699 0.06021 2.3064  0.001 *** | |  |  |  |  |  |
| Residual       108  27.6258 0.93979 | |  |  |  |  |  |
| Total          111  29.3957 1.00000 | |  |  |  |  |  |
| --- |  |  |  |  |  |  |
| Signif. codes:  0 ‘***’ 0.001 ‘**’ 0.01 ‘*’ 0.05 ‘.’ 0.1 ‘ ’ 1 | |  |  |  |  |  |
| permutest(betaTG) |  |  |  |  |  |  |
| Permutation test for homogeneity of multivariate dispersions | | |  |  |  |  |
| Permutation: free |  |  |  |  |  |  |
| Number of permutations: 999 |  |  |  |  |  |  |
| Response: Distances |  |  |  |  |  |  |
| Df  Sum Sq  Mean Sq     F N.Perm Pr(>F) | |  |  |  |  |  |
| Groups      3 0.11430 0.038100 8.067    999  0.001 *** | |  |  |  |  |  |
| Residuals 108 0.51008 0.004723 | |  |  |  |  |  |
| --- |  |  |  |  |  |  |
| Signif. codes:  0 ‘***’ 0.001 ‘**’ 0.01 ‘*’ 0.05 ‘.’ 0.1 ‘ ’ 1 | |  |  |  |  |  |
| **5b) wUnifrac** |  |  |  |  |  |  |
| Permutation test for adonis under reduced model | |  |  |  |  |  |
| Terms added sequentially (first to last) |  |  |  |  |  |  |
| Permutation: free |  |  |  |  |  |  |
| Number of permutations: 999 |  |  |  |  |  |  |
| adonis2(formula = DISTWAR ~ TreatmentGroup, data = FUCREA) | | |  |  |  |  |
| Df SumOfSqs      R2    F Pr(>F) | |  |  |  |  |  |
| TreatmentGroup   3  0.09352 0.08444 3.32  0.005 ** | |  |  |  |  |  |
| Residual       108  1.01406 0.91556 | |  |  |  |  |  |
| Total          111  1.10758 1.00000 | |  |  |  |  |  |
| --- |  |  |  |  |  |  |
| Signif. codes:  0 ‘***’ 0.001 ‘**’ 0.01 ‘*’ 0.05 ‘.’ 0.1 ‘ ’ 1 | |  |  |  |  |  |
| Permutation test for homogeneity of multivariate dispersions | | |  |  |  |  |
| Permutation: free |  |  |  |  |  |  |
| Number of permutations: 999 |  |  |  |  |  |  |
| Response: Distances |  |  |  |  |  |  |
| Df  Sum Sq   Mean Sq      F N.Perm Pr(>F) | |  |  |  |  |  |
| Groups      3 0.00761 0.0025366 1.2512    999  0.288 | |  |  |  |  |  |
| Residuals 108 0.21896 0.0020274 | |  |  |  |  |  |
| **5c) Bray Reproductive State** |  |  |  |  |  |  |
| adonis2(DISTWAR ~ ReproState, data = FUCREA) | |  |  |  |  |  |
| Permutation test for adonis under reduced model | |  |  |  |  |  |
| Terms added sequentially (first to last) |  |  |  |  |  |  |
| Permutation: free |  |  |  |  |  |  |
| Number of permutations: 999 |  |  |  |  |  |  |
| adonis2(formula = DISTWAR ~ ReproState, data = FUCREA) | | |  |  |  |  |
| Df SumOfSqs      R2      F Pr(>F) |  |  |  |  |  |  |
| ReproState   1   0.5866 0.01995 2.2397  0.005 ** | |  |  |  |  |  |
| Residual   110  28.8092 0.98005 | |  |  |  |  |  |
| Total      111  29.3957 1.00000 |  |  |  |  |  |  |
| Permutation test for homogeneity of multivariate dispersions | | |  |  |  |  |
| Permutation: free |  |  |  |  |  |  |
| Number of permutations: 999 |  |  |  |  |  |  |
| Response: Distances |  |  |  |  |  |  |
| Df  Sum Sq  Mean Sq      F N.Perm Pr(>F) | |  |  |  |  |  |
| Groups      1 0.09332 0.093322 19.284    999  0.001 *** | |  |  |  |  |  |
| Residuals 110 0.53233 0.004839 | |  |  |  |  |  |
| --- |  |  |  |  |  |  |
| Signif. codes:  0 ‘***’ 0.001 ‘**’ 0.01 ‘*’ 0.05 ‘.’ 0.1 ‘ ’ 1 | |  |  |  |  |  |
| **5d) wUnifrac  Reproductive State** |  |  |  |  |  |  |
| Permutation test for adonis under reduced model | |  |  |  |  |  |
| Terms added sequentially (first to last) |  |  |  |  |  |  |
| Permutation: free |  |  |  |  |  |  |
| Number of permutations: 999 |  |  |  |  |  |  |
| adonis2(formula = DISTWAR ~ ReproState, data = FUCREA) | | |  |  |  |  |
| Df SumOfSqs      R2      F Pr(>F) |  |  |  |  |  |  |
| ReproState   1  0.06166 0.05567 6.4853  0.007 ** | |  |  |  |  |  |
| Residual   110  1.04592 0.94433 | |  |  |  |  |  |
| Total      111  1.10758 1.00000 |  |  |  |  |  |  |
| --- |  |  |  |  |  |  |
| Signif. codes:  0 ‘***’ 0.001 ‘**’ 0.01 ‘*’ 0.05 ‘.’ 0.1 ‘ ’ 1 | |  |  |  |  |  |
| Permutation test for homogeneity of multivariate dispersions | | |  |  |  |  |
| Permutation: free |  |  |  |  |  |  |
| Number of permutations: 999 |  |  |  |  |  |  |
| Response: Distances |  |  |  |  |  |  |
| Df  Sum Sq   Mean Sq      F N.Perm Pr(>F) | |  |  |  |  |  |
| Groups      3 0.00761 0.0025366 1.2512    999  0.309 | |  |  |  |  |  |
| Residuals 108 0.21896 0.0020274 | |  |  |  |  |  |
| **5e) Bray Animal ID** |  |  |  |  |  |  |
| Permutation test for homogeneity of multivariate dispersions | | |  |  |  |  |
| Permutation: free |  |  |  |  |  |  |
| Number of permutations: 999 |  |  |  |  |  |  |
| Response: Distances |  |  |  |  |  |  |
| Df  Sum Sq  Mean Sq      F N.Perm Pr(>F) | |  |  |  |  |  |
| Groups    31 0.48203 0.015549 1.1547    999  0.282 | |  |  |  |  |  |
| Residuals 80 1.07726 0.013466 | |  |  |  |  |  |
| Lm-Exposure |  |  |  |  |  |  |
| Permutation test for homogeneity of multivariate dispersions | | |  |  |  |  |
| Permutation: free |  |  |  |  |  |  |
| Number of permutations: 999 |  |  |  |  |  |  |
| Response: Distances |  |  |  |  |  |  |
| Df  Sum Sq  Mean Sq      F N.Perm Pr(>F) | |  |  |  |  |  |
| Groups    31 0.48203 0.015549 1.1547    999  0.304 | |  |  |  |  |  |
| Residuals 80 1.07726 0.013466 |  |  |  |  |  |  |
| **5f) wUnifrac Animal ID** |  |  |  |  |  |  |
| Permutation test for adonis under reduced model | |  |  |  |  |  |
| Terms added sequentially (first to last) |  |  |  |  |  |  |
| Permutation: free |  |  |  |  |  |  |
| Number of permutations: 999 |  |  |  |  |  |  |
| adonis2(formula = DISTWAR ~ SequecningID, data = FUCREA) | | |  |  |  |  |
| Df SumOfSqs      R2      F Pr(>F) |  |  |  |  |  |  |
| SequecningID  7  0.33758 0.38982 1.9166  0.002 ** | |  |  |  |  |  |
| Residual     21  0.52841 0.61018 | |  |  |  |  |  |
| Total        28  0.86599 1.00000 |  |  |  |  |  |  |
| --- |  |  |  |  |  |  |
| Signif. codes:  0 ‘***’ 0.001 ‘**’ 0.01 ‘*’ 0.05 ‘.’ 0.1 ‘ ’ 1 | |  |  |  |  |  |
| Permutation test for homogeneity of multivariate dispersions | | |  |  |  |  |
| Permutation: free |  |  |  |  |  |  |
| Number of permutations: 999 |  |  |  |  |  |  |
| Response: Distances |  |  |  |  |  |  |
| Df   Sum Sq   Mean Sq      F N.Perm Pr(>F) | |  |  |  |  |  |
| Groups     7 0.041651 0.0059501 1.5701    999  0.202 | |  |  |  |  |  |
| Residuals 21 0.079585 0.0037897 | |  |  |  |  |  |
| **5g) Bray APO** |  |  |  |  |  |  |
| Permutation test for adonis under reduced model | |  |  |  |  |  |
| Terms added sequentially (first to last) |  |  |  |  |  |  |
| Permutation: free |  |  |  |  |  |  |
| Number of permutations: 999 |  |  |  |  |  |  |
| adonis2(formula = DISTWAR ~ APO, data = FUCREA) | |  |  |  |  |  |
| Df SumOfSqs      R2      F Pr(>F) |  |  |  |  |  |  |
| APO       1   0.4605 0.06602 1.9086  0.018 * | |  |  |  |  |  |
| Residual 27   6.5150 0.93398 |  |  |  |  |  |  |
| Total    28   6.9755 1.00000 |  |  |  |  |  |  |
| --- |  |  |  |  |  |  |
| Signif. codes:  0 ‘***’ 0.001 ‘**’ 0.01 ‘*’ 0.05 ‘.’ 0.1 ‘ ’ 1 | |  |  |  |  |  |
| Permutation test for homogeneity of multivariate dispersions | | |  |  |  |  |
| Permutation: free |  |  |  |  |  |  |
| Number of permutations: 999 |  |  |  |  |  |  |
| Response: Distances |  |  |  |  |  |  |
| Df   Sum Sq   Mean Sq      F N.Perm Pr(>F) | |  |  |  |  |  |
| Groups     1 0.000046 0.0000458 0.0105    999  0.931 | |  |  |  |  |  |
| Residuals 27 0.117539 0.0043533 |  |  |  |  |  |  |
| **5h) wUnifrac APO** |  |  |  |  |  |  |
| Permutation test for adonis under reduced model | |  |  |  |  |  |
| Terms added sequentially (first to last) |  |  |  |  |  |  |
| Permutation: free |  |  |  |  |  |  |
| Number of permutations: 999 |  |  |  |  |  |  |
| adonis2(formula = DISTWAR ~ APO, data = FUCREA) | |  |  |  |  |  |
| Df SumOfSqs      R2      F Pr(>F) |  |  |  |  |  |  |
| APO       1  0.04733 0.05466 1.5611  0.139 | |  |  |  |  |  |
| Residual 27  0.81865 0.94534 |  |  |  |  |  |  |
| Total    28  0.86599 1.00000 |  |  |  |  |  |  |
| Permutation test for homogeneity of multivariate dispersions | | |  |  |  |  |
| Permutation: free |  |  |  |  |  |  |
| Number of permutations: 999 |  |  |  |  |  |  |
| Response: Distances |  |  |  |  |  |  |
| Df   Sum Sq   Mean Sq      F N.Perm Pr(>F) | |  |  |  |  |  |
| Groups     1 0.000041 0.0000410 0.0113    999  0.928 | |  |  |  |  |  |
| Residuals 27 0.098123 0.0036342 | |  |  |  |  |  |
| **Fig 7 Differential Abundance Statistics** |  |  |  |  |  |  |
| **7a)** |  |  |  |  |  |  |
| Genus | baseMean | log2FoldChange | p-value | Median abundance Non-pregnant | Median abundance pregnant | |
| RF39 | 243.151774 | 1.444696 | 6.06E-05 | 6.18E-03 | 1.17E-02 |  |
| Alloprevotella | 204.827994 | -2.246444 | 3.50E-09 | 6.78E-03 | 1.87E-03 |  |
| Terrisporobacter | 19.180944 | -4.6247 | 3.01E-06 | 7.29E-05 | 0.00E+00 |  |
| Rodentibacter | 10.808046 | -2.281721 | 8.33E-05 | 2.43E-04 | 4.50E-05 |  |
| Actinobacillus | 6.00085 | -3.121656 | 9.64E-05 | 7.95E-05 | 0.00E+00 |  |
| Romboutsia | 3.545022 | -3.386751 | 1.09E-04 | 1.14E-04 | 0.00E+00 |  |
| Bilophila | 3.190994 | 3.536582 | 3.52E-05 | 0.00E+00 | 0.00E+00 |  |
| Agathobacter | 1.872939 | -3.050138 | 7.99E-05 | 6.18E-05 | 0.00E+00 |  |
| **7b)** |  |  |  |  |  |  |
| Genus | baseMean | log2FoldChange | p-value | Median abundance No | Median abundance Yes | |
| Blautia | 168.40168 | 2.017203 | 1.03E-04 | 0.001663671 | 0.00678376 |  |
| Helicobacter | 72.66462 | 8.463221 | 5.81E-06 | 0 | 0.00135552 |  |
| Prevotellaceae_UCG-004 | 42.18705 | 8.52556 | 1.32E-04 | 0 | 0.00015061 |  |
| Akkermansia | 19.60936 | 21.251195 | 8.04E-07 | 0 | 0 |  |
| Lachnospiraceae_XPB1014_group | 13.46508 | 19.658367 | 8.71E-08 | 0 | 0 |  |
| **7c)** |  |  |  |  |  |  |
| Genus | baseMean | log2FoldChange | p-value | Median abundance No | Median abundance Yes | |
| Methanobrevibacter | 396.696878 | 2.995007 | 1.50E-08 | 2.36E-03 | 1.75E-02 |  |
| Streptococcus | 377.694482 | 4.615353 | 1.21E-04 | 2.70E-04 | 5.78E-03 |  |
| Marvinbryantia | 82.615352 | 1.888051 | 3.58E-04 | 1.03E-03 | 3.25E-03 |  |
| Mogibacterium | 74.659619 | 2.51554 | 6.77E-06 | 7.48E-04 | 2.90E-03 |  |
| p-1088-a5_gut_group | 47.892003 | 1.988268 | 1.05E-05 | 7.32E-04 | 2.71E-03 |  |
| Intestinibacter | 24.968 | 2.715043 | 5.03E-04 | 1.34E-04 | 5.87E-04 |  |
| Enterorhabdus | 23.451519 | 1.82075 | 7.30E-05 | 3.40E-04 | 9.68E-04 |  |
| Akkermansia | 19.609358 | 20.513371 | 5.61E-10 | 0.00E+00 | 0.00E+00 |  |
| Lachnospira | 11.990607 | -4.174677 | 3.15E-05 | 2.83E-04 | 4.01E-05 |  |
| Eggerthellaceae uncultured_bacterium | 11.439181 | 1.993898 | 1.16E-04 | 1.21E-04 | 6.16E-04 |  |
| unclassified, Order: Oscillospirales | 11.063787 | 4.04988 | 1.15E-04 | 8.83E-06 | 1.85E-04 |  |
| Incertae_Sedis | 8.809012 | 2.204371 | 1.51E-04 | 8.20E-05 | 3.70E-04 |  |
| **7d)** |  |  |  |  |  |  |
| Genus | baseMean | log2FoldChange | p-value | Median abundance No | Median abundance Yes | |
| Akkermansia | 136.662 | 12.60141 | 5.47E-06 | 0 | 0 |  |
